# RUNX1-deficiency drives immune-active ER^+^ mammary tumorigenesis through activation of interferon signaling

**DOI:** 10.64898/2026.04.06.716728

**Authors:** Sen Han, Dongxi Xiang, Xueqing Chen, Dongyi Zhao, Guyu Qin, Roderick T. Bronson, Zhe Li

## Abstract

Recurrent *loss-of-function* mutations in *RUNX1* occur in estrogen receptor-positive (ER^+^) breast cancers, yet how RUNX1-loss contributes to breast tumorigenesis remains unclear. Here we used genetically engineered mouse models with luminal mammary epithelial cell (MEC)-restricted gene disruption to investigate its role in breast cancer initiation. Loss of RUNX1 alone, or together with RB1, was insufficient to drive tumor formation. In contrast, combined loss of RUNX1 and p53 induced mammary tumors with full penetrance. These tumors contained ER^+^ cancer cells and exhibited extensive T cell and macrophage infiltration, indicative of an immune hot microenvironment. Mechanistically, RUNX1-deficiency activated interferon signaling in luminal MECs, associated with derepression of RUNX1 target *STAT1* and enhanced inflammatory responses. Consistent with these findings, human ER^+^ breast cancers with low *RUNX1* expression displayed elevated immune signatures and poorer patient survival. Together, our results identify RUNX1-loss as a driver of an immune-active subtype of ER^+^ breast cancer.

## INTRODUCTION

High throughput sequencing-based genomic studies have generated a comprehensive catalog of mutations in human breast cancers ^1–4^. Critical next steps following these efforts are to identify and validate driver mutations and to understand how they contribute to breast tumorigenesis. Many of the significantly mutated genes identified in these studies are already known to play roles in breast cancer when altered, such as *TP53*, *PIK3CA*, *GATA3* and *CDH1*. In addition, several novel recurrent mutations affecting transcription factors (TFs) or transcriptional regulators have emerged, such as *RUNX1* (and its co-factor gene *CBFB*), *TBX3*, *FOXA1*, and *CTCF* ^4^. Among them, *RUNX1* and *CBFB* are of particular interest.

RUNX1 (Runt-related transcription factor 1), together with RUNX2 and RUNX3, and their common non-DNA-binding, obligate partner CBFβ, constitute a small family of TFs ^5^. All three RUNX/CBFβ TFs play critical roles in both normal development and disease. RUNX1 is an essential regulator of hematopoiesis and immunity ^5,6^. Mutations or translocations involving *RUNX1* and *CBFB* are frequently observed in leukemia ^5^. In addition, loss of RUNX1 function has been implicated in autoimmune disease (e.g., psoriasis, rheumatoid arthritis, and systemic lupus erythematosus) and inflammation ^7–10^. RUNX2 is a master regulator of osteoblast differentiation and bone development ^11^, whereas RUNX3 plays important roles in cytotoxic T-cell development as well as in gastrointestinal and neural development ^12^. More recently, the roles of RUNX TFs in epithelial cancers, particularly breast cancer, have begun to emerge ^13^.

Mutations of *RUNX1* and *CBFB* in human breast cancer are restricted to estrogen receptor-positive (ER^+^) tumors and are predicted to be *loss-of-function* ^4,14^, suggesting that RUNX1 may function as a tumor suppressor in ER^+^ breast cancer. In our previous work ^14^, we demonstrated that RUNX1 acts as a master regulator of ER^+^ luminal mammary epithelial cells (MECs). Specifically, we found that RUNX1 represses *Elf5*, a master regulatory TF gene for alveolar luminal cells, and regulates mature luminal TF/co-factor genes (e.g., *Foxa1* and *Cited1*) involved in the ER transcriptional program ^14^. In addition, RUNX1 was shown to antagonize estrogen-mediated suppression of AXIN1, and its loss in ER^+^ luminal breast cancer cells leads to increased levels of centrosomal phospho-β-catenin and mitotic deregulation, which may contribute to hyperproliferation of RUNX1-deficient ER^+^ luminal cells ^15^. RUNX1 has also been reported to stabilize the MEC phenotype by preventing epithelial-to-mesenchymal transition (EMT) and to suppress breast cancer stem cell activity and breast tumor growth ^16–18^. However, despite these findings, a causal role of RUNX1-deficiency in breast tumorigenesis has not yet been established.

In our previous study, we showed that MEC-specific deletion of *Runx1* by *MMTV-Cre* impaired the fate of ER^+^ luminal cells, leading to a reduction in their population size. Co-deletion of *Trp53* (or *Rb1*) was able to rescue *Runx1*-null ER^+^ luminal MECs, resulting in their hyperproliferation ^14^. However, *MMTV-Cre*-mediated loss of *Runx1* together with *Trp53* (or *Rb1*) caused early lethality in mice, likely due to leaky expression of *MMTV-Cre* in hematopoietic cells and the subsequent development of hematopoietic malignancies. Consequently, we were unable to determine whether the rescued *Runx1*-null ER^+^ luminal MECs could progress to mammary tumor. Here, we report a novel mouse model that strictly restricts Cre-mediated deletion of *Runx1* and *Trp53* (or *Rb1*) to MECs. Using this model, we demonstrate that loss of RUNX1 cooperates with p53-deficiency to drive the development of immune-active ER^+^ mammary tumors with complete penetrance. Our findings provide new insights into how RUNX1-deficiency contributes to the development of a distinct subset of ER^+^ breast cancer characterized by increased immune “hotness”.

## RESULTS

### RUNX1-deficiency cooperates with p53-loss, but not with RB1-loss, to promote mammary tumor development from luminal cells

Since *MMTV-Cre*-driven mice did not allow us to determine whether *Runx1/Trp53*-null or *Runx1/Rb1*-null MECs could progress to mammary tumors, we used a breast cancer induction approach we established based on intraductal injection of a Keratin 8 (K8)^+^ luminal MEC-specific Cre-expressing adenovirus (*Ad-K8-Cre*)^19^ into the mammary glands of female mice carrying conditional knockout alleles of *Runx1*, *Trp53*, or *Rb1*, as well as a conditional Cre-reporter, *Rosa26-LSL-YFP* (*R26Y*) ^20^. The cohorts included *Runx1^L/L^;Trp53^L/L^;R26Y* (*RxPY*) and *Runx1^L/L^;Rb1^L/L^;R26Y* (*RRY*) mice, as well as control strains *Rb1^L/L^;Trp53^L/L^;R26Y* (*RbPY*), *Trp53^L/L^;R26Y* (*PY*), *Runx1^L/L^;R26Y* (*RxY*), and *R26Y*-only mice. Following intraductal *Ad-K8-Cre* injection, we performed lineage-tracing of YFP-marked MECs in injected mammary glands using Fluorescence-Activated Cell Sorting (FACS) analysis. Significant expansion of YFP^+^ MECs over time was observed in all mice carrying the *Trp53* conditional knockout alleles (i.e., *RxPY*, *RbPY*, and *PY* mice, Fig. 1a and Supplementary Fig. 1a). In contrast, no significant expansion of YFP^+^ cells was detected in the injected *RRY* mice, although a small number of YFP-marked *Runx1/Rb1*-null (as well as *Runx1*-null or WT) cells persisted long-term in injected glands (Fig. 1a and Supplementary Fig. 1a).

**Fig. 1.**
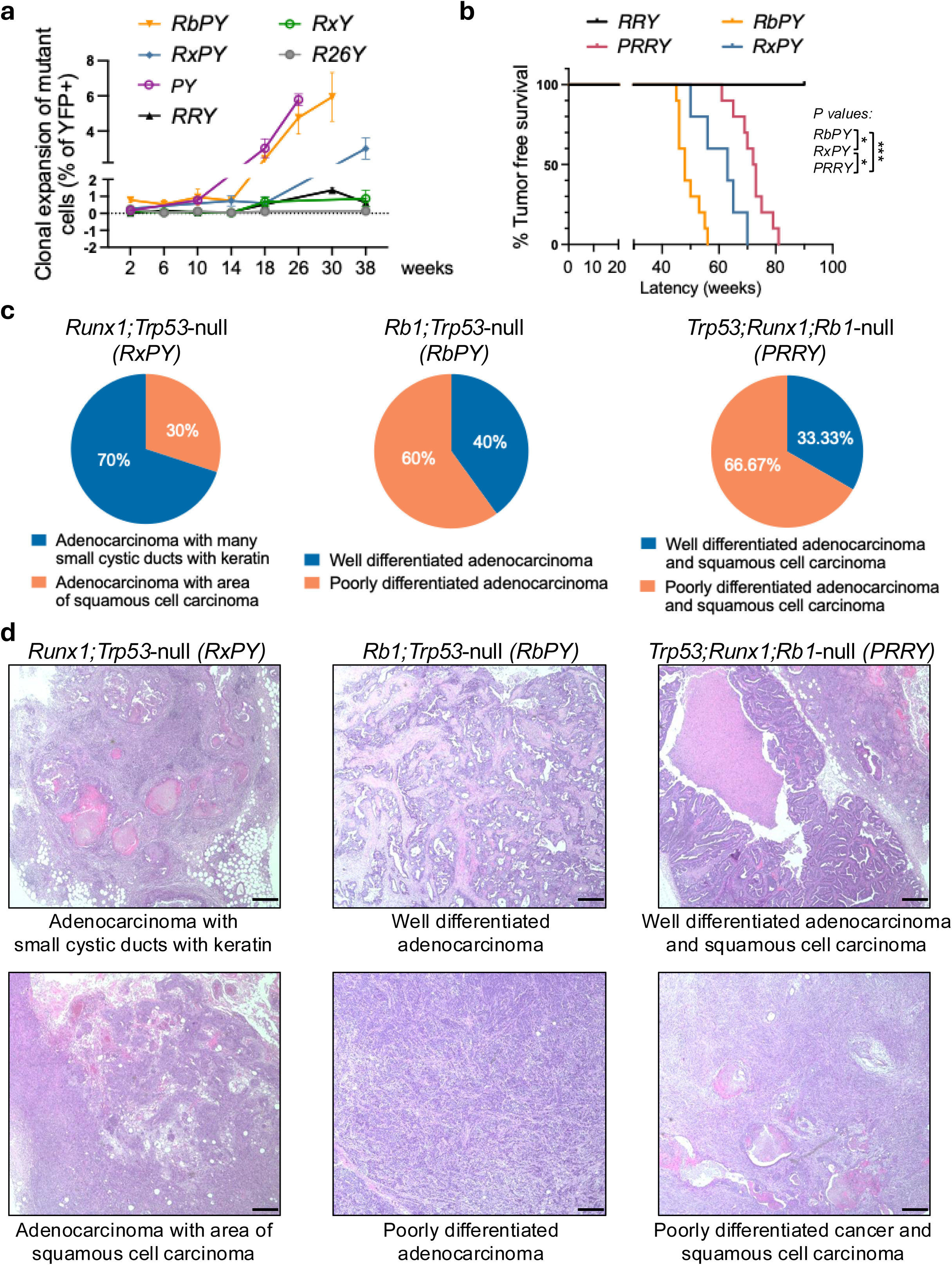
Characterization of mutant MECs and mammary tumorigenesis in the indicated models. Cre recombination was induced via intraductal *Ad-K8-Cre* injection in 8-week-old female mice carrying the indicated conditional knockout alleles and the *R26Y* reporter. (a) Expansion of mutant or WT (*R26Y*) luminal MECs (as % of YFP^+^ cells within the lineage-negative compartment) was assessed by FACS analysis starting from 2 weeks post-induction. There were at least 3 replicates at each time point per mouse model (n≥3). (b) Kaplan-Meier tumor free curve for each model was established once the mammary tumor became palpable. Each group contains 10 mice (n=10). (c) Mammary tumors observed from cohorts in b were further designated into different categories based on histological examination results; at least 8 slides were assessed for each type of tumor. (d) Representative Hematoxylin and Eosin (H&E) staining pictures for different histologic types of tumors as in c. Scale bar = 50μm.

Since *Ad-K8-Cre*-mediated recombination restricts gene deletion to MECs in the mammary gland, this system allowed us to determine whether mutant MECs eventually progress to mammary tumors. Consistent with the clonal expansion of *Runx1/Trp53*-null MECs, all injected *RxPY* mice developed mammary tumors after a relatively long latency (Fig. 1b). In contrast, none of the similarly injected *RRY* female mice developed tumors (Fig. 1b). In the injected *RbPY* control female mice, clonal expansion of *Rb1/Trp53*-null YFP^+^ premalignant MECs was followed by mammary tumor development in all injected mice, with a shorter latency than observed in *RxPY* mice (Fig. 1b).

We reported previously that induced loss of p53 alone in luminal MECs of *PY* mice, achieved through intraductal *Ad-K8-Cre* injection, also resulted in mammary tumors with full penetrance, and that tumors arising in this model were predominantly Claudin-low-like poorly differentiated cancers, often accompanied by focal amplification of a genomic region containing the proto-oncogene *Yap1* ^21^. In contrast, mammary tumors arising in the *Ad-K8-Cre*-injected *RxPY* mice exhibited different histological features. Histological analysis revealed that almost all *RxPY* tumors were adenocarcinomas mixed with either small cystic ducts containing keratin pearls or areas of squamous cell carcinoma (Fig. 1c-d). These tumors also differed from those arising in similarly induced *RbPY* control mice, which primarily developed either well- or poorly differentiated adenocarcinomas lacking keratin deposition or squamous metaplasia (Fig. 1c-d).

Since *Ad-K8-Cre*-injected *RRY* mice failed to develop tumors, we next asked whether additional loss of p53 would enable mammary tumor formation. To test this, we generated *Trp53^L/L^;Runx1^L/L^;Rb1^L/L^;R26Y* (*PRRY*) mice. Following intraductal *Ad-K8-Cre* injection, all *PRRY* developed mammary tumors with complete penetrance, although with a longer latency than observed in *RxPY* mice (Fig. 1b). Histological analysis showed that *PRRY* tumors resembled *RxPY* tumors more closely than *RbPY* tumors: most were well- or poorly differentiated adenocarcinomas containing areas of squamous cell carcinoma (Fig. 1c-d).

The presence of squamous metaplasia in most *RxPY* and *PRRY* tumors suggests activation of Wnt/β-catenin signaling, a pathway previously linked to this phenotype ^22,23^. Consistent with this, recent *in vivo* imaging studies have shown that constitutive activation of Wnt/β-catenin signaling can induce squamous transdifferentiation of MECs ^24^. Interestingly, RUNX1-loss has been reported to promote aberrant activation of Wnt/β-catenin signaling in epithelial cells ^15,25^. To determine whether this pathway is activated in *Runx1*-null tumors, we performed immunostaining for β-catenin and LEF1 in *RxPY* and *PRRY* tumors, using *RbPY* tumors as controls. *Runx1*-null tumors, particularly *RxPY* tumors, displayed stronger β-catenin staining than *RbPY* tumors and importantly, many *Runx1*-null tumor cells showed nuclear β-catenin staining that overlapped with LEF1 staining, whereas *RbPY* tumor cells predominantly exhibited membrane-localized β-catenin (Supplementary Fig. 1b).

Together, these findings indicate that *Runx1*-loss and *Rb1*-loss independently cooperate with p53-deficiency to drive mammary tumorigenesis, giving rise to distinct tumor phenotypes. We found no evidence that *Runx1*-loss directly cooperates with *Rb1*-loss in luminal MECs to promote mammary tumor development. The characteristic squamous metaplasia observed in *Runx1*-null tumors is likely attributable to aberrant activation of Wnt/β-catenin signaling.

### RUNX1-deficiency cooperates with p53-loss to drive development of ER^+^ mammary tumors

To determine the MEC types represented in tumors arising from these different genetic models, we performed FACS analysis using a previously established gating strategy that separates Lin^⁻^CD24^hi^CD29^lo^ (Lin: lineage markers) luminal MECs into Sca-1^+^ and Sca-1^⁻^ subsets, corresponding to ER^+^ and ER^⁻^ luminal MECs, respectively ^26^ (Supplementary Fig. 2). In *RxPY* tumors, the majority of Lin⁻CD24^hi^CD29^lo^ luminal cells were Sca-1^+^, and the proportion of Sca-1^+^ cells increased further when gating specifically on YFP^+^ luminal cells (Fig. 2a-b). A similar trend was observed in *PRRY* tumors, although with greater variability: in most *PRRY* tumors, 80-100% luminal cells were Sca-1^+^, whereas one tumor contained <50% Sca-1^+^ luminal cells (Fig. 2a-b). In contrast, *RbPY* tumors generally contained fewer Sca-1^+^ cells than tumors from the *Runx1*-deficient models (Fig. 2a-b). Most *RbPY* tumors had ≤ 50% of Sca-1^+^ luminal cells, even when analysis was restricted to YFP^+^ luminal cells (Fig. 2a-b).

**Fig. 2.**
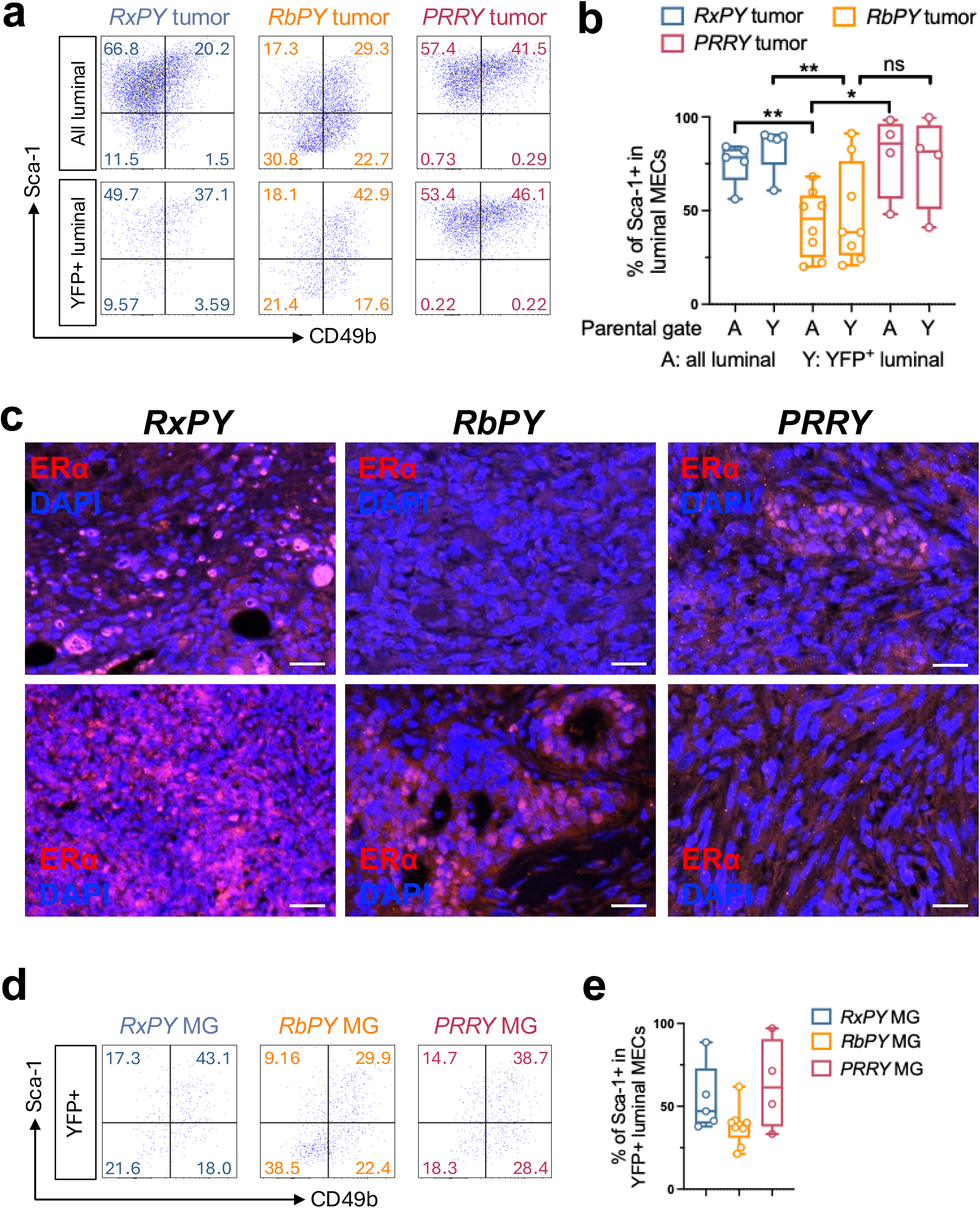
RUNX1-loss facilitates ER^+^ mammary tumor development. (a) Representative FACS plots showing expression of Sca-1 and CD49b in either all tumor cells within the luminal gate (lineage⁻CD24^hi^CD29^+^, upper panel) or YFP-marked tumor cells within the luminal gate (lineage⁻YFP^+^CD24^hi^CD29^+^, lower panel) based on the parental gating shown in Supplementary Fig. 2. (b) Statistical analysis of Sca-1^+^ cells based on the FACS analysis shown in a was performed on *RxPY* (n=5)*, RbPY* (n=8), and *PRRY* (n=4) tumors. (c) Immunofluorescence staining showing ERα^+^ cells (red, nuclear staining) in *RxPY, RbPY,* and *PRRY* tumors. Scale bar = 25μm. (d, e) FACS analysis for Sca-1 and CD49b expression in YFP^+^ mutant luminal MECs from *RxPY* (n=5)*, RbPY* (n=8), and *PRRY* (n=4) premalignant mammary glands (6 months after induction).

To validate these FACS results, we performed immunostaining for ERα in tumor sections from each model. Nearly all *RxPY* tumors, including both well- and poorly differentiated adenocarcinomas, contained regions with abundant ERα^+^ tumor cells (Fig. 2c, left), consistent with their Sca-1 positivity (Fig. 2b). In contrast, most *RbPY* tumors, particularly poorly differentiated adenocarcinomas, stained negative for ERα, although a small subset of more differentiated *RbPY* tumors with glandular morphology exhibited strong ERα positivity (Fig. 2c, middle); this is in line with the FACS data showing that in a small of such tumors, most YFP^+^ luminal cells were Sca-1^+^ (Fig. 2b). In the *PRRY* model, most tumors also contained areas enriched for ERα^+^ tumor cells, whereas a small number of poorly differentiated tumors were largely ERα-negative (Fig. 2c, right), also in line with the FACS data for Sca-1 (Fig. 2b). Together, these findings indicate that, consistent with the association of *RUNX1* mutations with ER^+^ human breast cancers, *Runx1*-loss in murine luminal MECs cooperates with p53-deficiency to drive the development of ER^+^ mammary tumors.

To determine whether the ERα^+^ tumor cells observed in these models may originate from ER^+^ luminal MECs, we further analyzed YFP-marked premalignant MECs by FACS using the same Sca-1-based gating strategy (Supplementary Fig. 2). On average, close to 50% of YFP^+^ premalignant luminal MECs in induced *RxPY* mice were Sca-1^+^, and this proportion was slightly higher in *PRRY* mice (Fig. 2d-e). In contrast, most induced *RbPY* mice contained <50% of Sca-1^⁺^ YFP^+^ premalignant luminal MECs (Fig. 2d-e). These findings support the possibility that *Runx1*-null mammary tumors, particularly those arising in the *RxPY* model, originate from ER^+^ luminal MECs that undergo clonal expansion during the premalignant stage.

### Molecular characterization of *RxPY*, *RbPY*, and *PRRY* mammary tumors

To further characterize mammary tumors arising in *Ad-K8-Cre*-induced *RxPY*, *RbPY*, and *PRRY* mice at the molecular level, we performed RNA sequencing (RNA-seq). Since mammary tumors from *PRRY* mice exhibited histological features similar to those from *RxPY* mice, we grouped these tumors together as *Runx1*-null tumors and compared them with *Rb1*-null tumors from *RbPY* mice using gene-set enrichment analysis (GSEA) ^27^.

All tumors analyzed in this study were generated in the context of p53-loss. We reported previously that induced loss of p53 alone in luminal MECs leads to the development of Claudin-low-like mammary tumors, often accompanied by focal amplification of a genomic region containing the proto-oncogene *Yap1*, and less frequently another region spanning *Met* ^21^. To evaluate whether similar genomic amplifications were present in our profiled tumors, we used GSEA-based expression analysis as a surrogate for genomic amplification by examining elevated expression of genes within these regions. This analysis identified two tumors (PRb-6 and PRR-1) with elevated expression of most genes in the *Yap1* locus, suggesting amplification of this region; PRR-1 also showed potential amplification of the *Met* locus (Supplementary Fig. 3a). To determine their tumor subtype identities, we compared their expression profiles with those from a large panel of murine mammary tumor models^28^ and with human breast cancer intrinsic subtypes. These two tumors showed the highest similarity to the murine and human Claudin-low subtype (Supplementary Fig. 3b-c, highlighted). Since the phenotype of these tumors is likely dominated by amplification of the proto-oncogene *Yap1*, we excluded them from subsequent comparisons between *Runx1*-null and *Rb1*-null tumors.

GSEA analysis revealed that *Runx1*-null tumors most closely resembled the Squamous_like^Ex^ murine subtype (Fig. 3a), consistent with the frequent squamous metaplasia observed histologically in these tumors (Fig. 1c-d). *Runx1*-null tumors also exhibited similarities to the Claudin-low^Ex^, Class8^Ex^, and Stat1^Ex^ murine subtypes (Fig. 3a). Of note, the Stat1^Ex^ subtype, derived from the *Stat1*^⁻/⁻^ mouse model, is characterized by the development of ERα^+^ mammary tumors ^28,29^. In contrast, *Rb1*-null tumors showed the highest similarity to the C3Tag^Ex^, Neu^EX^, and Myc^EX^ murine subtypes (Fig. 3b). When compared with human breast cancer intrinsic subtypes, *Runx1*-null tumors most closely resembled the Claudin-low and Normal-like subtypes, whereas *Rb1*-null tumors were most similar to the Luminal B subtype (Fig. 3c-d).

**Fig. 3.**
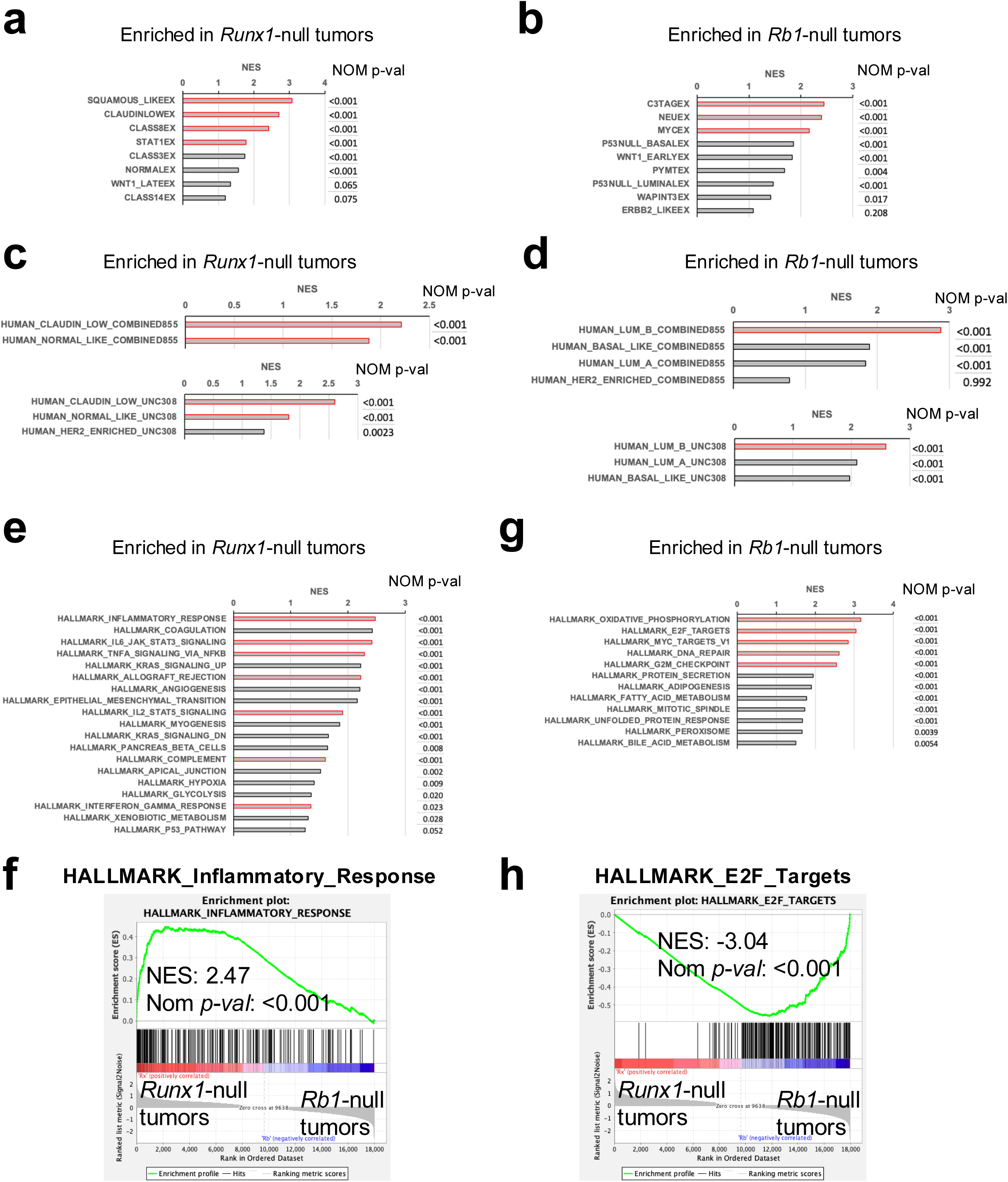
RNA-seq analysis of *RxPY*, *RbPY,* and *PRRY* mammary tumors. (a-d) GSEA data showing top enriched gene sets representing mouse mammary tumor (a,b) or human breast cancer (c,d) intrinsic subtypes extracted from Pfefferle et al (human intrinsic subtypes were based on the UNC308 data set and Combined855 data set)^28^ in either *Runx1*-null tumors (*RxPY* and *PRRY*) (a,c) or *Rb1*-null tumors (*RbPY*) (b,d). (e-h) GSEA data showing top enriched gene sets in the Hallmark collection from the MSigDB in either *Runx1*-null tumors (*RxPY* and *PRRY*) (e) or *Rb1*-null tumors (*RbPY*) (g). Gene sets related to inflammation/immune pathways (e) or E2F/cell cycle (g) are highlighted, and representative GSEA enrichment plots are shown in f and h.

We next compared *Runx1*-null and *Rb1*-null tumors at the pathway level using GSEA with the Hallmark gene set collection. In *Runx1*-null tumors, the most significantly enriched gene sets were associated with inflammation, angiogenesis, and EMT, whereas *Rb1*-null tumors were enriched for gene sets related to cell proliferation, including oxidative phosphorylation, E2F targets, and MYC targets (Fig. 3e-h, highlighted). As RB1 functions primarily as a tumor suppressor by binding and inhibiting E2F transcription factors, loss of RB1 is expected to result in activation of E2Fs and E2F target genes, consistent with the observed transcriptional profile of *Rb1*-null tumors.

To validate these findings at the cellular level, we stained *RxPY*, *RbPY*, and *PRRY* tumors for Ki67, a marker of cell proliferation. Most *RbPY* tumors contain abundant Ki67^+^ cells, whereas *RxPY* tumors exhibited relatively few Ki67^+^ cells, and *PRRY* tumors showed intermediate levels (Supplementary Fig. 3d). The upregulation of inflammation-related gene sets in *Runx1*-null tumors is of particular interest, as it suggests that loss of RUNX1 in luminal MECs may trigger an innate immune response, leading to increased immune activity within the tumor microenvironment. These transcriptional differences may help explain the distinct tumor latencies observed in these models. The *RbPY* model exhibits the shortest latency, likely reflecting the higher proliferation rate of *Rb1*-null premalignant and tumor cells (Fig. 1a and Supplementary Fig. 3d). In contrast, the *RxPY* and *PRRY* models show longer tumor latencies, potentially due to RUNX1-loss-induced inflammation and immune reactions, which may slow tumor progression.

### RUNX1-deficient mammary tumors exhibit extensive immune cell infiltrations

To validate the immune-related transcriptional signatures observed in *Runx1*-null mammary tumors, we performed multi-color FACS analysis of tumors from *RxPY*, *RbPY*, and *PRRY* mice (Supplementary Fig. 4). Both RUNX1-deficient *RxPY* and *PRRY* tumors contained significantly higher proportions of CD45^+^ leukocytes than RUNX1-WT *RbPY* tumors, with several *PRRY* tumors exhibiting the highest levels of tumor-infiltrating immune cells (Fig. 4a-b).

**Fig. 4.**
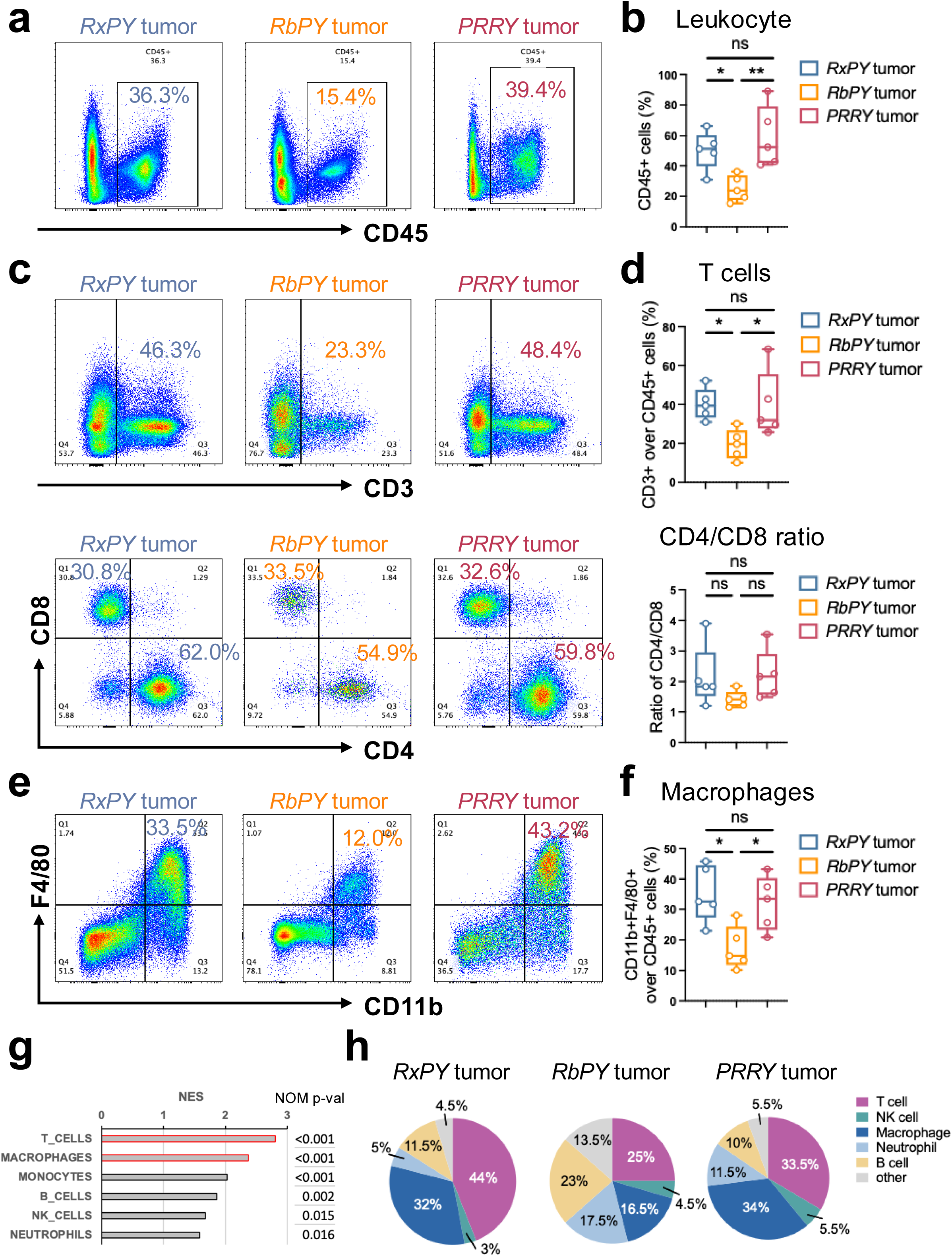
Enhanced immune “hotness” in *Runx1*-null mammary tumors. Profiling of immune cell populations within the microenvironment of *RxPY* (n=5)*, RbPY* (n=5), and *PRRY* (n=5) tumors by FACS and their corresponding statistical analysis. (a, b) The overall immune cell infiltration was presented by the proportion of CD45^+^ cells in the tumor single cell suspension (dead cells excluded from the parental gate). (c, d) Tumor-infiltrating T cells were characterized by CD3^+^ cells within the CD45^+^ compartment (upper panel). CD4^+^ T and CD8^+^ T cells were further evaluated within the CD45^+^CD3^+^ gate (lower panel). (e, f) Tumor-associated macrophages were identified by CD11b^+^F4/80^+^ within the CD45^+^ gate. In a-f, representative FACS plots for each tumor type are shown at left (a,c,e) and summarized data are shown at right (b,d,f). (g) GSEA of tumor RNA-seq data showing top-enriched immune cell signatures (highlighted) in *Runx1*-null tumors in relation to *Rb1*-null tumors; gene sets representing different immune cell population signatures are derived from Nirmal et al ^30^. (h) Pie charts represent the average proportions of T cells (CD45^+^CD3^+^), NK cells (CD45^+^CD3^-^NKp46^+^), macrophages (CD45^+^CD11b^+^F4/80^+^), neutrophils (CD45^+^CD11b^+^Ly-6G^+^), and B cells (CD45^+^CD19^+^) within the indicated tumor types (n=5 each). *P* value: \**p*<0.05, \*\**p*<0.01, ns = not significant. Data represent mean ± S.E.M.

We next examined the composition of immune cell subpopulations within these tumors. The increased immune infiltration observed in *RxPY* and *PRRY* tumors was largely attributable to elevated numbers of CD3^+^ T cells (Fig. 4c-d, top plots) and F4/80^+^CD11b^+^ macrophages (Fig. 4e-f). Within the CD3^+^ T cell compartment, analysis of CD4^+^ and CD8^+^ T cell subsets revealed slightly higher CD4/CD8 ratios in *RxPY* and *PRRY* tumors compared with *RbPY* tumors, although this difference did not reach statistical significance (Fig. 4c-d, bottom plots). Importantly, as *RxPY* and *PRRY* tumors contained higher overall proportions of CD3^+^ T cells and CD45^+^ leukocytes, these tumors should likely harbor greater absolute numbers of CD8^+^ T cells than *RbPY* tumors.

Since the gene expression profiles shown in Fig. 3 were generated from whole tumors, which include contributions from both tumor cells and infiltrating immune cells, we further assessed enrichment of immune cell-specific gene expression signatures^30^ in these datasets. Consistent with our FACS results, signatures corresponding to T cells and macrophages were the most strongly enriched immune cell signatures in *Runx1*-null tumors (Fig. 4g).

Lastly, we examined additional major immune cell populations within the tumors. Compared with *Rb1*-null *RbPY* tumors, *Runx1*-null *RxPY* and *PRRY* tumors contained lower proportions of neutrophils and B cells (Fig. 4h). Together, these results demonstrate that RUNX1-loss is associated with increased immune cell filtration in mammary tumors, particularly involving T cells and macrophages, consistent with the inflammatory gene expression signatures observed in *Runx1*-null tumors.

### Induced loss of RUNX1 in p53-deficient luminal cells leads to increased T cell infiltration

To determine whether the increased immune cell infiltration observed in *Runx1*-null tumors results primarily from RUNX1-loss in MECs rather than from secondary genetic or epigenetic changes acquired during tumor progression, we analyzed immune cell populations in intraductally injected mammary glands at the premalignant stage by FACS analysis (Supplementary Fig. 5). Compared with glands with induced loss of RB1 (and p53), glands with induced loss of RUNX1 (and p53) exhibited an almost 2-fold increase in the infiltrating CD45^+^ leukocytes (Fig. 5a-b). Further analysis of immune cell subsets revealed that this increase was largely driven by a significant expansion of CD3^+^ T cell population, which constituted more than half of the immune cells in injected *RxPY* mammary glands (Fig. 5c-d, top plots).

**Fig. 5.**
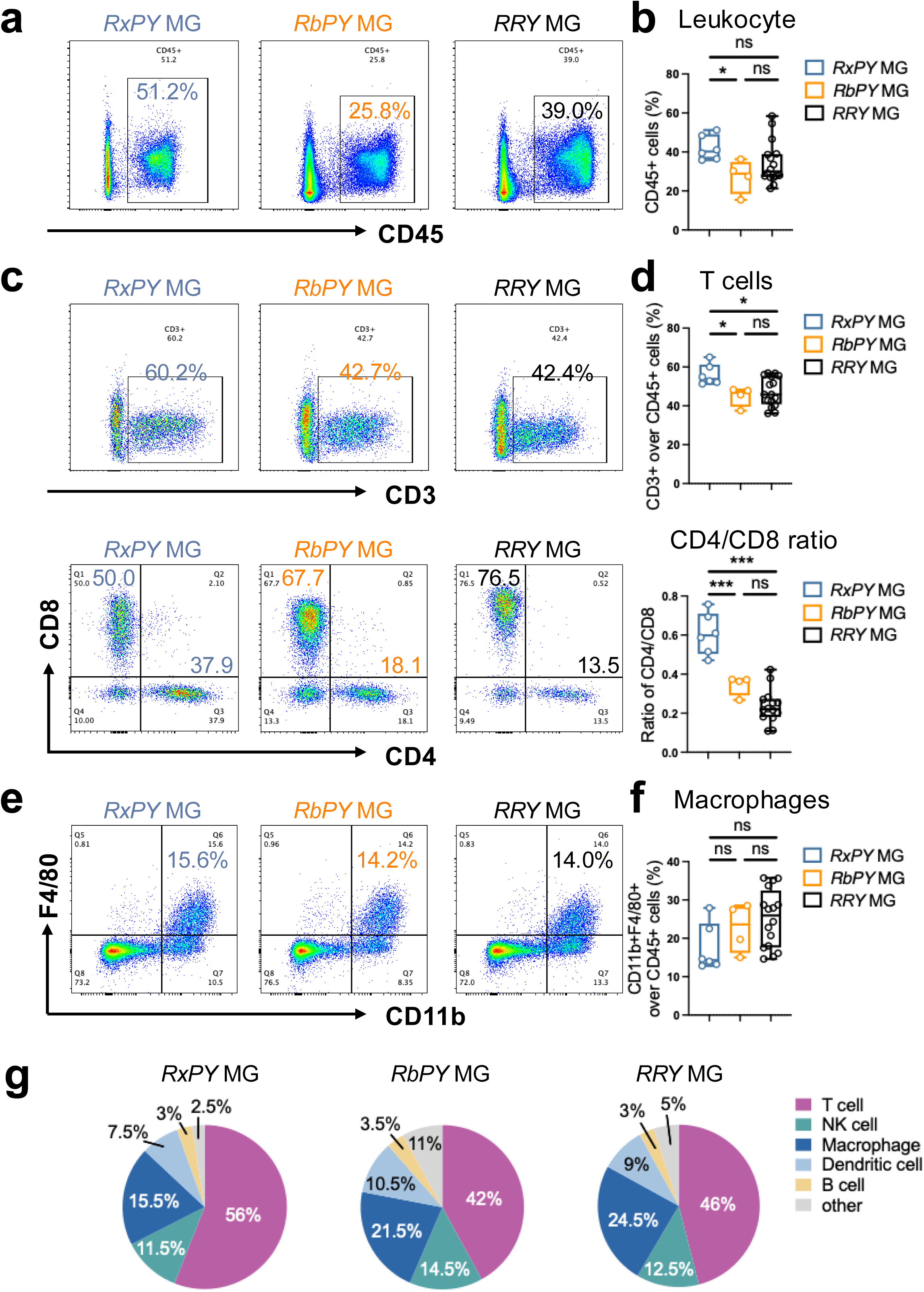
Loss of RUNX1 in p53-deficient luminal MECs results in increased T cell infiltration. Profiling of overall immune cells (a, b), T cells and T cell subsets (c, d), and macrophages (e, f) within the premalignant mammary glands (all around 6 months of chasing) of *RxPY* (n=6)*, RbPY* (n=4), and *RRY* (n=16) mice by FACS (representative FACS plots are shown in a,c,e, corresponding statistical analyses are shown in b,d,f). (g) Pie charts represent the average proportions of T cells (CD3^+^), NK cells (CD3^-^NKp46^+^), macrophages (CD11b^+^F4/80^+^), dendritic cells (CD11b^+^CD11c^+^), and B cells (CD19^+^) within the CD45 gate in premalignant mammary glands. *P* value: \**p*<0.05, \*\**p*<0.01, ns = not significant. Data represent mean ± S.E.M.

Within the mammary gland CD3^+^ T cell compartment, more than half of the cells were CD8^+^ T cells (Fig. 5c-d, bottom plots). This distribution differs from that observed in the resulting mammary tumors, where CD4^+^ T cells predominated (Fig. 4c-d, bottom plots). This shift may reflect increased expansion of regulatory CD4^+^ T cells and/or a reduction in cytotoxic CD8^+^ T cells during tumor progression. Of note, although the percentage of CD8^+^ T cells in the injected *RxPY* glands was somewhat lower than in the injected *RbPY* or *RRY* control glands, this was apparently due to a higher proportion of CD4^+^ T cells (i.e., higher CD4/CD8 ratio, Fig. 5d, bottom). This trend persisted in the resulting *Runx1*-null tumors, with *RxPY* tumors generally exhibiting higher CD4/CD8 ratios than *RbPY* tumors, although this difference did not reach statistical significance (Fig. 4d, bottom). Importantly, despite the lower percentage of CD8^+^ T cells in the injected *RxPY* glands, the absolute number of CD8^+^ T cells is still expected to be higher than in the injected *RbPY* glands, owing to the overall increase in CD3^+^ T cells and CD45^+^ leukocytes in *RxPY* glands (Fig. 5a-d).

We also examined the myeloid compartment and found that the injected *RxPY* and *RbPY* mammary glands contained similar proportions of F4/80^+^CD11b^+^ macrophages (Fig. 5e-f). Of note, in the *RxPY* model, this macrophage population expanded markedly in the resulting mammary tumors (Fig. 4e-f) compared with premalignant mammary glands (Fig. 5e-f), indicating a potential role of macrophages during progression of *Runx1*-null tumors. Among other major immune cell populations, we did not detect significant differences between *RxPY* and *RbPY* glands (Fig. 5g). Together, these findings suggest that induced loss of RUNX1 in luminal MECs triggers an early immune response, characterized by initial infiltration of CD8^+^ T cells, followed by expansion of CD4^+^ T cells and, at later stages, substantial accumulation of macrophages in the developing mammary tumors.

### Loss of RUNX1 in p53-deficient luminal cells activates interferon signaling

Since induced loss of RUNX1 in p53-deficient luminal MECs resulted in early T cell infiltration in the mammary gland microenvironment, we reasoned that the increased immune “hotness” observed in *Runx1*-null mammary tumors might primarily arise from RUNX1-loss in MECs themselves. To investigate the molecular basis for this phenomenon, we analyzed YFP-marked *Runx1/Trp53*-null premalignant luminal MECs (2-weeks after induced loss of RUNX1) and compared them with similarly generated *Trp53*-null only and WT luminal MECs following intraductal *Ad-K8-Cre* injection. Gene expression profiling was performed and analyzed using GSEA. This revealed that, relative to either WT (Supplementary Fig. 6a) or *Trp53*-null MECs (Supplementary Fig. 6b), the most significantly enriched Hallmark gene sets in *Runx1/Trp53*-null MECs were associated with cell cycle and proliferation, while the second most enriched category comprised interferon (IFN) response-related gene sets (Fig. 6a-b and Supplementary Fig. 6a-b). Importantly, the enrichment of IFN-related signatures was not attributable to loss of p53 alone in MECs, as these signatures remained elevated when *Runx1/Trp53*-null MECs were compared directly with *Trp53*-null MECs. Nor was it due to adenoviral injection, as *R26Y*-only and *PY* control mice underwent the same procedure. These findings suggest that the increased immune/IFN signature may result from the loss of transcriptional control by RUNX1.

**Fig. 6.**
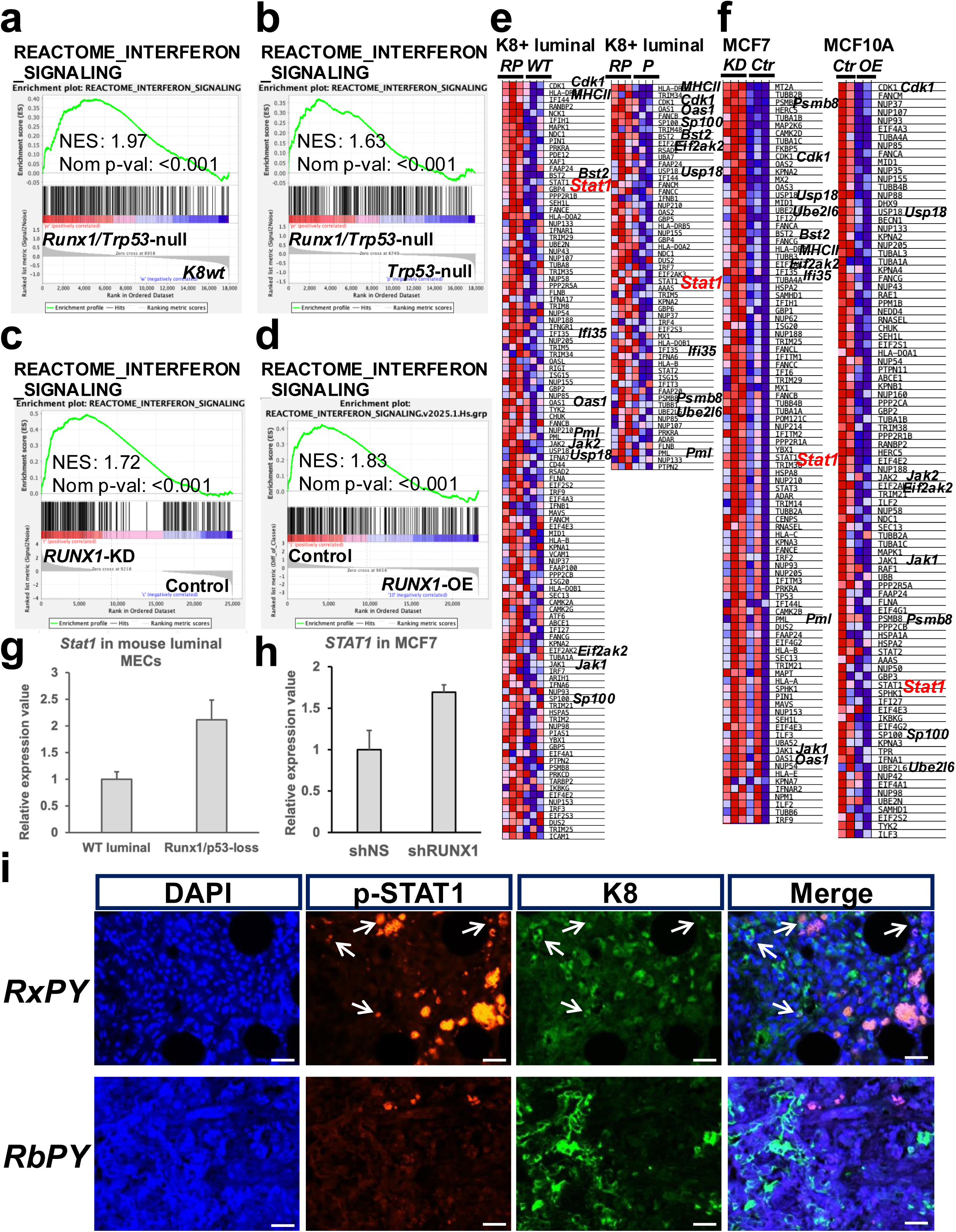
IFN response pathway is active in RUNX1-deficient ER^+^ luminal cells. (a-d) GSEA plots showing significant enrichment of various IFN pathway gene sets in mouse *Runx1/Trp53*-null luminal MECs compared to WT (a) or *Trp53*-null (b) controls, as well as in human MCF7 breast cancer cells with *RUNX1* knockdown (KD) compared to its control (c) and in MCF10A control cells compared to MCF10A cells with *RUNX1* overexpression (OE) (d). (e-f) Heatmaps corresponding to GSEA plots in a-d; key genes upregulated in mouse *Runx1/Trp53*-null luminal cells, human MCF7 cells with *RUNX1*-KD, and human MCF10A control cells (compared to *RUNX1* OE) are indicated. In the heatmaps, red to blue represent the highest to lowest expression. *RP*: *Runx1/Trp53*-null; *P*: *Trp53*-null; *Ctr*: control. (g-h) Upregulation of *STAT1* expression in mouse *Runx1/Trp53-null* luminal cells (compared to WT controls, g) and in MCF7 cells with *RUNX1* KD (based on GEO #GSE75070) (shRUNX1, h). (i) Co-immunofluorescence staining showing K8 and phosphorylated STAT1 (p-STAT1) double positive cells (arrows) in *RxPY* tumor, but not in *RbPY* tumor. Scale bars = 25μm.

To further explore this possibility, we compared gene expression profiles of *Runx1/Trp53*-null premalignant luminal MECs with those of similarly generated *Brca1/Trp53*-null premalignant MECs, which eventually give rise to BRCA1-deficient basal-like tumors ^31^. GSEA showed that immune/IFN-related gene sets were the most significantly enriched Hallmark signatures in *Runx1/Trp53*-null cells relative to *Brca1/Trp53*-null cells (Supplementary Fig. 6c). As *Brca1/Trp53*-null premalignant MECs accumulate DNA damage ^31^, which can also induce inflammation but does so over time, the rapid enrichment of IFN-related signatures in *Runx1/Trp53*-null premalignant MECs only 2-weeks after induction suggests a direct transcriptional mechanism rather than a secondary response to genomic instability and DNA damage is the more likely cause.

We next examined publicly available expression datasets from human breast cells with either *RUNX1* knockdown (KD) ^32^ or overexpression (OE) ^18^. In the ER^+^ breast cancer cell line MCF7, *RUNX1* KD resulted in significant upregulation of IFN-related gene sets (Fig. 6c and Supplementary Fig. 6d). Conversely, in MCF10A cells, an immortalized luminal progenitor-like normal human breast epithelial cell line, *RUNX1* OE led to downregulation of multiple IFN-related gene sets, which appeared enriched in vector-only control samples (Fig. 6d). Importantly, many IFN pathway genes altered by *RUNX1* loss or OE were shared between mouse and human luminal MECs (Fig. 6e-f). Since these IFN-related transcriptional changes were observed in cultured cells lacking immune components, together with the findings from our mouse model, these data support a MEC-intrinsic mechanism (e.g., related to RUNX1 transcriptional control) linking RUNX1-loss to IFN pathway activation, at least during the early stages.

Several mechanisms could explain how RUNX1 regulates IFN signaling. One possibility is that IFN response genes are direct transcriptional targets of RUNX1 and are normally repressed by it. Alternatively, RUNX1 may regulate (i.e., repress) a master regulatory TF controlling IFN signaling. To investigate these possibilities, we analyzed RUNX1-binding peaks in MCF7 cells from a publicly available Chromatin Immunoprecipitation-Sequencing (ChIP-seq) dataset ^32^. Although RUNX1 binding was not detected in most IFN response genes, we identified three RUNX1-binding peaks within the *STAT1* gene locus: one in the promoter region (proximal), one upstream (distal) site, and one within a downstream intron (intronic) (Supplementary Fig. 6e). Interestingly, the proximal and distal binding sites lie within narrow valleys of H3K27Ac (acetylation of lysine 27 residue in histone H3) signal, a histone modification associated with active chromatin. This pattern suggests that RUNX1 binding at these sites may function to repress *STAT1* transcription.

Consistent with this interpretation, *Stat1* expression was increased in both *Runx1/Trp53*-null premalignant mouse luminal MECs and *RUNX1*-KD human MCF7 cells, relative to their respective controls (Fig. 6e-f: heatmaps, g-h: relative levels). Analysis of human breast cancer datasets further revealed a negative correlation between *RUNX1* and *STAT1* expression, particularly in ER^+^ luminal A breast cancers (Supplementary Fig. 6f). Moreover, clinical outcome analyses indicated that higher *STAT1* expression correlates with poorer prognosis in ER^+^ breast cancers or the luminal A breast cancer subtype, whereas lower *STAT1* expression correlates with worse outcomes in ER^⁻^ breast cancers or the basal-like breast cancer subtype (Supplementary Fig. 6g). Intriguingly, *RUNX1* shows the opposite pattern of association with clinical outcomes (Supplementary Fig. 6h), suggesting both RUNX1 and STAT1 may play distinct and opposing roles depending on breast cancer subtype and ER status.

To confirm activation of the IFN signaling pathway in *Runx1*-null mammary tumors, we performed co-immunostaining for phospho-STAT1 (pSTAT1) and K8. STAT1 is a key TF in IFN signaling that activates type I and II IFN response genes upon phosphorylation by JAK1, JAK2 or TYK2 ^33^. We observed that *RxPY* tumors frequently contained pSTAT1^+^ cells, some of which co-expressed K8, indicating epithelial tumor cells (Fig. 6i, arrows). Additional pSTAT1^+^ K8^⁻^cells with fibroblast-like appearance were also present, likely representing activated (inflamed) stromal cells or tumor cells that have undergone EMT. In contrast, *RbPY* tumors contained markedly fewer pSTAT1^+^ cells (Fig. 6i).

Together, these findings indicate that RUNX1-loss in luminal MECs induces activation of IFN signaling, at least in part through derepression (thus upregulation) of the RUNX1 target gene *STAT1*. Elevated STAT1 levels may enhance its interaction with IFN receptors and activation by them ^34^, ultimately promoting transcription of IFN response genes.

### Human RUNX1-deficient ER^+^ breast tumors exhibit increased immune/IFN signatures

To evaluate the relevance of findings from our *Runx1*-null murine models to human disease, we analyzed human breast tumors from the METABRIC dataset, one of the largest publicly available human breast cancer genomic cohorts ^35,36^. Since *RUNX1* mutations occur at relatively low frequency, and *RUNX1* expression may also be reduced through mechanisms other than somatic mutation ^15^, we defined a statistically more meaningful subset of RUNX1-deficient human ER^+^ breast cancers based on *RUNX1* transcript levels. Within the METABRIC cohort, ER^+^ tumors were stratified into *RUNX1*-low and *RUNX1*-high groups, corresponding to approximately the lowest and highest 10% of *RUNX1* expression, respectively (Fig. 7a). Importantly, tumors in these two groups exhibited similar levels of *ESR1* expression (Fig. 7a).

**Fig. 7.**
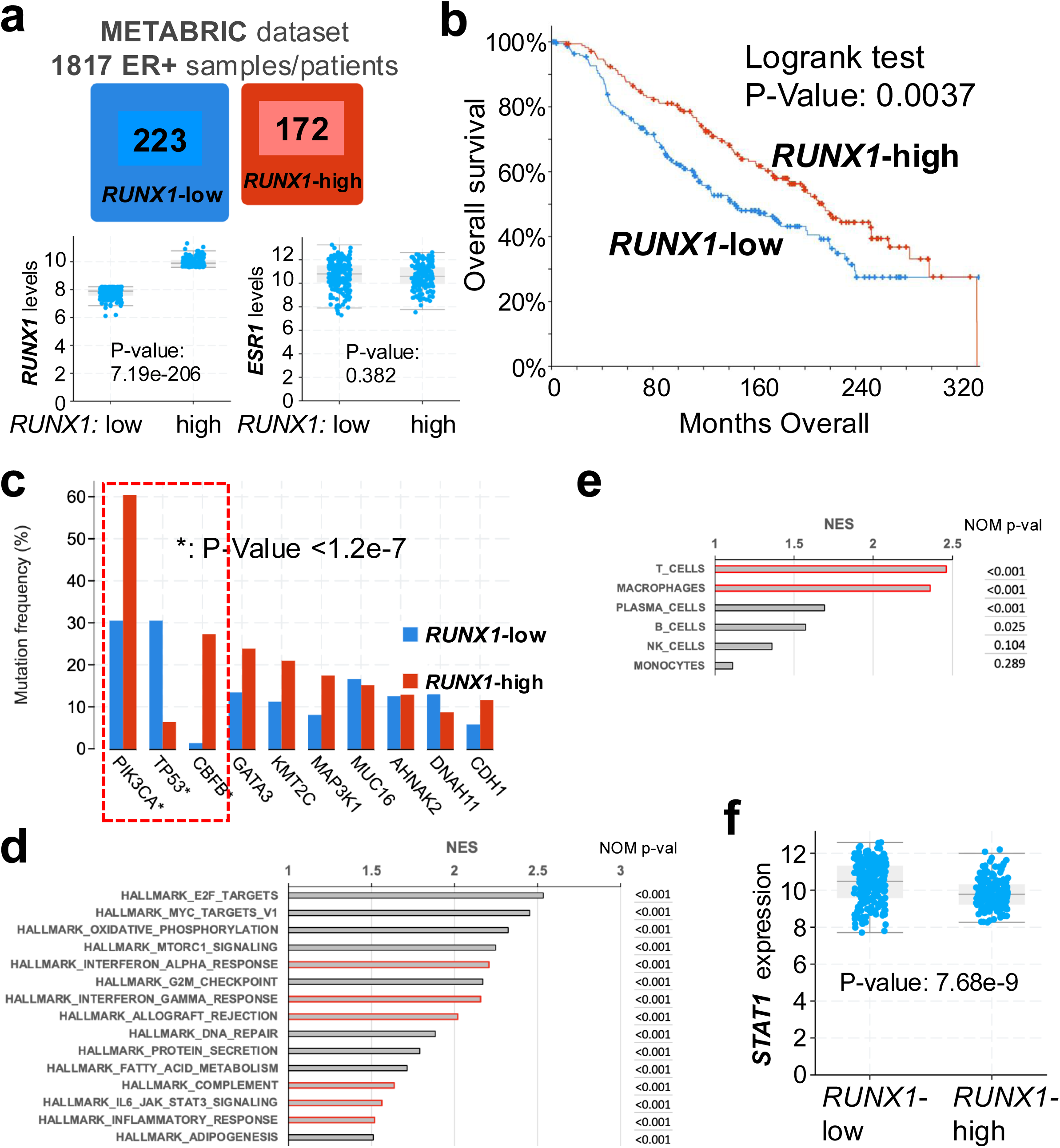
Defining a *RUNX1*-low subset of ER^+^ breast tumors from the METABRIC cohort (cbioportal). (a) 223 and 172 cases from 1817 ER^+^ breast tumor samples in the METABRIC cohort with the lowest and highest levels of *RUNX1* (see lower left plot) are assigned to the *RUNX1*-low and *RUNX1*-high groups, respectively; these cases have similar levels of *ESR1* expression (lower right plot). (b) Patient survival analysis showing *RUNX1*-low cases exhibited significantly worse overall survival. (c) Other mutations associated with *RUNX1*-low vs. *RUNX1*-high groups; significant association is marked by *. (d) GSEA data showing top enriched gene sets in the Hallmark collection from the MSigDB in *RUNX1*-low tumors (compared to *RUNX1*-high tumors); immune-related gene sets are highlighted. (e) GSEA showing top-enriched immune cell signatures (highlighted) in *RUNX1*-low tumors (compared to *RUNX1*-high tumors); gene sets representing different immune cell population signatures are derived from Nirmal et al ^30^. (f) Significantly higher levels of *STAT1* expression in *RUNX1*-low tumors (compared to *RUNX1*-high tumors).

Kaplan-Meier survival analysis showed that patients with *RUNX1*-low tumors had significantly worse overall survival compared with those with *RUNX1*-high tumors (Fig. 7b). Mutation analysis further revealed that although most somatic mutations were similarly distributed between the two groups, *TP53* mutations were strongly enriched in *RUNX1*-low tumors (Fig. 7c). This observation is consistent with our experimental findings that p53-loss rescues RUNX1-deficient ER^+^ MECs ^14^ and with our development of a RUNX1-deficient tumor model requiring combined loss of p53 and RUNX1. In contrast, *CBFB* (loss-of-function) mutations are significantly associated with the *RUNX1*-high group (Fig. 7c). This mutually exclusive relationship likely reflects the functional dependency of RUNX1 on its cofactor CBFβ, encoded by *CBFB*, such that *CBFB loss-of-function* mutations can phenocopy RUNX1-deficiency ^5^.

We next compared gene expression profiles between *RUNX1*-low and *RUNX1*-high tumors. Consistent with findings from our *Runx1*-null premalignant MECs and mouse tumors, gene sets associated with IFN signaling and inflammatory pathways were among the most strongly enriched Hallmark gene sets in *RUNX1*-low tumors (Fig. 7d, highlighted). We further examined enrichment of immune cell-specific expression signatures^30^ and found that signatures corresponding to T cells and macrophages were the two most strongly enriched immune populations in *RUNX1*-low tumors (Fig. 7e). This pattern closely parallels the immune infiltration observed in *Runx1*-null mammary tumors in our mouse models (Fig. 4). We also assessed expression of the RUNX1 target gene *STAT1* in these tumors and found that *STAT1* expression was significantly higher in *RUNX1*-low tumors compared with *RUNX1*-high tumors (Fig. 7f).

To validate these observations in an independent dataset, we performed a similar analysis using the TCGA breast cancer cohort ^4^. ER^+^ tumors were again stratified into *RUNX1*-low and *RUNX1*-high groups, and consistent with the METABRIC results, *RUNX1*-low tumors displayed stronger immune and IFN-related gene signatures, indicative of increased tumor immune “hotness” (Supplementary Fig. 7a-f).

Together, these analyses of human breast cancer datasets support the key conclusion from our murine studies: loss of RUNX1 in breast epithelial cells is associated with activation of IFN signaling and the development of immune hot breast cancers.

## DISCUSSION

In this mouse modeling study, we showed that induced loss of RUNX1 alone, or combined loss of RUNX1 and RB1, in luminal MECs is insufficient to drive mammary tumorigenesis. In contrast, combined loss of RUNX1 and p53 leads to the development of a distinct class of mammary tumors with full penetrance. These tumors are characterized by a highly immune-active microenvironment with extensive infiltration of T cells and macrophages, the presence of ER^+^ tumor cells, and areas of squamous metaplasia. The latter phenotype likely results from aberrant activation of Wnt/β-catenin signaling in *Runx1*-null MECs, a previously described consequence of RUNX1-loss ^15,22–25^. Although human breast cancers with squamous metaplasia are rare and have not been reported to be associated with RUNX1-deficient ER^+^ breast cancers, the ability of RUNX1 to negatively regulate Wnt/β-catenin signaling appears conserved between mice and humans. The squamous metaplasia observed in our mouse models may therefore reflect species-specific responses to Wnt/β-catenin activation in MECs.

Although induced loss of p53 alone can drive mammary tumorigenesis, as we reported previously ^21^, tumors arising in the *Trp53*-null model are typically driven by secondary genomic alterations, such as amplification of the *Yap1* genomic region. In contrast, such amplification was not observed in most *Runx1*-null tumors in this study (Supplementary Fig. 3a). Moreover, tumors arising in the *RxPY* model (i.e., loss of both RUNX1 and p53) are histologically distinct from those in the *PY* model (i.e., loss of p53 alone). Together, these findings indicate that mammary tumor development in the *RxPY* model cannot be attributed solely to p53-loss and provide direct evidence that RUNX1-deficiency acts as a driver of a specific subtype of mammary tumors.

The identification of RUNX1-deficient tumors as immune hot tumors is particularly intriguing. RUNX1 is a key regulator of immune function, and its loss has been implicated in several autoimmune and inflammatory diseases, including psoriasis, rheumatoid arthritis, and systemic lupus erythematosus ^7–10^. Most previously described roles of RUNX1 in regulating immunity involve its functions in immune cells. In contrast, our findings demonstrate that in luminal MECs, RUNX1 not only maintains the ER^+^ luminal lineage ^14^, but also suppresses innate immune signaling pathways within epithelial cells. Our data suggest that RUNX1-loss in luminal MECs primarily activates IFN signaling, which may initially promote anti-tumor immune responses, such as recruitment of CD8^+^ cytotoxic T cells (Fig. 5c-d). This immune activation may explain the relatively long tumor latency observed in our RUNX1-deficient mouse models, as many cells in the resulting tumors remain pSTAT1^+^ (Fig. 6i), indicative of active IFN signaling. However, chronic immune activation within the tumor microenvironment may eventually promote tumor progression, potentially contributing to the poorer clinical outcomes observed in patients with RUNX1-low ER^+^ breast cancers (Fig. 7b and Supplementary Fig. 7b).

Multiple mechanisms may contribute to activation of IFN and inflammation pathways following RUNX1-loss in luminal MECs. Our analysis of premalignant MECs with induced RUNX1 loss, together with data from human breast cell lines, suggests that RUNX1-deficiency may activate IFN response genes at least in part through derepression of the RUNX1 target gene *STAT1*. Elevated STAT1 levels may enhance downstream signaling through its interaction with IFN receptors ^34^, ultimately promoting transcription of IFN-stimulated genes. In addition, immune activation may also emerge during later stages through co-evolution between RUNX1-deficient MECs and the immune microenvironment. These mechanisms are not mutually exclusive and may jointly contribute to the immune “hotness” in RUNX1-deficient tumors.

Interestingly, in RUNX1-deficient ER^+^ MECs, we also observed increased signatures of ribosome biogenesis, which may contribute to activation of IFN signaling as well. Emerging evidence indicates that ribosome biogenesis can play an important role in regulating innate immune response to virus infection and cytosolic DNA ^37^. For example, inhibition of ribosome biogenesis reduces production of IFNB1 and IFN-stimulated genes, thereby promoting viral replication ^37^. Conversely, enhanced ribosome biogenesis observed in patients with Hutchinson-Gilford progeria syndrome, a premature aging disease, is associated with increased IFN production ^38^. Hyperactive ribosome biogenesis can destabilize rDNA loci due to conflicts between rDNA transcription and replication, leading to DNA double-strand breaks ^39^ and activation of the DNA damage response ^40^. Such perturbations may promote accumulation of cytosolic DNA and activation of DNA-sensing pathways, including the cGAS-STING pathway, ultimately leading to activation of IFN signaling.

Mutations in *RUNX1* occur predominantly in ER^+^ breast cancer. Although ER^+^ breast cancers generally have a favorable prognosis, they account for ∼70-80% of all breast cancer cases ^41,42^, are highly heterogeneous, and remain associated with significant morbidity and mortality. Treatment failure often results from resistance to endocrine therapy or, in metastatic disease, resistance to CDK4/6 inhibitors ^43^. Our findings suggest that RUNX1-deficient ER^+^ breast cancers represent a distinct subset characterized by worse overall survival and increased immune activity.

Compared with triple-negative breast cancers (TNBCs) and HER2^+^ breast tumors, ER^+^ tumors typically contain fewer tumor-infiltrating leukocytes (TILs) ^44,45^, possibly reflecting the immunomodulatory effects of estrogen/ER signaling ^46,47^. Nonetheless, ER^+^ breast cancers exhibit substantial inter- and intra-tumoral heterogeneity in immune infiltration ^44,45,48–50^. As ER^+^ tumors generally display lower antigenicity ^45^, it remains unclear how these cancers recruit different levels of TILs and what their prognostic significance may be. Interestingly, high expression of an IFN metagene (i.e., IFN-induced genes) has been shown to predict distant metastasis specifically in ER^+^HER2^⁻^ tumors ^51^, suggesting that IFN signaling may play a functional role in certain ER^+^ breast cancers. Epidemiological studies further show that regular use of nonsteroidal anti-inflammatory drugs (NSAIDs, e.g., aspirin) reduces the risk of ER^+^ but not ER^⁻^ breast cancers ^52^, implying that ER^+^ tumors may be particularly responsive to inflammatory signaling and that inflammatory cytokines may promote more aggressive disease phenotypes ^46^. As ER^+^ breast cancer cells typically exhibit lower antigenicity, immunotherapies that enhance the activity of tumor-infiltrating lymphocytes, such as immune checkpoint blockade, may hold particular promise when appropriately targeted to immune-active subsets ^45^. A better understanding of how TILs are induced in ER^+^ breast cancer, and how tumor-intrinsic alterations such as RUNX1-loss shape the immune microenvironment, may therefore have important implications for patient stratification and immunotherapy strategies.

Overall, improved understanding of RUNX1-deficient breast cancers may reveal new therapeutic opportunities. Such insights could help overcome resistance to endocrine therapy and/or CDK4/6 inhibitors and may provide a rationale for combining targeted therapies with immune checkpoint blockade in this distinct subset of ER^+^ breast cancer.

## METHODS

### Mice

All mouse studies were approved under protocol 2020N000122 and conducted in compliance with the Institutional Animal Care and Use Committee (IACUC) guidelines at Brigham and Women’s Hospital. *Trp53^L/L^* mice (B6.129P2-*Trp53tm1^Brn^*/J, strain# 008462), *Rb1^L/L^*mice (B6;129-*Rb1^tm3Tyj^*/J, strain# 008186), and *Rosa26-stop-YFP* (*R26Y*) mice (B6.129X1-*Gt(ROSA)26Sor*^tm1(EYFP)Cos^/J, strain# 006148) were obtained from The Jackson Laboratory.

*Runx1^L/L^* mice (B6;129-*Runx1^tm3.1Spe^*/J, strain# 010673) were kindly provided by Dr. Alan Cantor (Boston Children’s Hospital). These mouse lines were intercrossed accordingly to obtain homozygous mouse lines, namely, *Trp53^L/L^*;*R26Y^homo^*(*PY*), *Runx1^L/L^;R26Y^homo^ (RxY), Runx1^L/L^;Rb1^L/L^;R26Y^homo^* (*RRY*), *Runx1^L/L^;Trp53^L/L^;R26Y^homo^* (*RxPY*), *Rb1^L/L^;Trp53^L/L^;R26Y^homo^* (*RbPY*), and *Trp53^L/L^*;*Runx1^L/L^; Rb1^L/L^;R26Y^homo^* (*PRRY*). Homozygous *R26Y^homo^* and *PY* mouse lines were backcrossed for at least six generations to FVB/NJ background, while homozygous *RxY, RbY, RxY, RRY*, and *PRRY* mouse lines were maintained on a mixed background of C57BL/6J and FVB/NJ.

### Tumor induction, lineage tracing, and sample preparation

The spontaneous breast cancer modeling approach has been described previously ^53^. Briefly, lineage tracing was initiated in female mice at 8-weeks of age. The *Keratin 8* (*K8*) promoter-driven Cre-expressing adenovirus (*Ad-K8-Cre*, VVC-Li-535, University of Iowa Viral Vector Core) was delivered into lumen through intraductal injection in order to induce Cre-mediated recombination in luminal mammary epithelial cells (MECs). Mice were analyzed at defined time points during the premalignancy. Tumor-bearing mice were euthanized prior to reaching humane endpoints, as determined by the animal protocol. Euthanasia was performed by carbon dioxide (CO_2_) overdose in Euthanex multi-cage chamber units, which were set to introduce 100% CO_2_ at a fill rate of 30% displacement of the chamber volume per minute with CO_2_, added to the existing air in the chamber.

Mammary tumor or premalignant (abdominal) mammary gland tissues were collected from mice subjected to lineage tracing. Each mammary tumor specimen was subdivided for downstream applications: one portion was snap-frozen in dry ice and stored at -80°C for subsequent bulk RNA sequencing; a second portion was fixed in 4% formaldehyde overnight and submitted to the Rodent Histopathology Core at Harvard Medical School for paraffin embedding and histological examination; the remaining portion was processed immediately for single-cell preparation.

For single-cell preparation, excised tumor fragments or premalignant mammary gland tissues were finely minced and digested in advanced DMEM/F12 supplemented with 2% fetal bovine serum (FBS), 1% HEPES, collagenase I (16.5 μg/5 mL), and DNase I (75 μg/5 mL) at 37°C for 1 hour with continuous agitation. Following digestion, samples were treated with red blood cell lysis buffer for 5 minutes at room temperature, washed, and filtered to obtain single-cell suspensions for follow-up analyses.

### Flow cytometry (FACS)

Single-cell suspensions of tumor or premalignant mammary gland tissues were freshly prepared from mice undergoing lineage tracing and were subjected to FACS analysis subsequently. Cells were first labeled with a fixable viability dye (1:500) for 20 minutes at room temperature in the dark. Fc receptor blocking was performed using anti-CD16/32 antibody (1:200; BioLegend, clone S17011E) at 4°C for 5 minutes. Surface staining was carried out with fluorophore-conjugated antibodies (see Supplementary Table 1) at 4°C for 30 minutes according to the manufactures’ protocols. Data acquisition was performed on BD LSR II or BD Symphony A5 instruments (BD Biosciences), and analyses were conducted by FloJo software (v10.10.0). Detailed gating strategies are provided in Supplementary Fig. 2, 4, and 5.

### Expression profiling and data analysis

For premalignant (or WT) luminal MEC samples, total RNAs from YFP^+^ MECs sorted from mammary glands of *PY* or *RxPY* or *Trp53^L/L^;Brca1^L/L^;R26Y* female mice ∼10-14 days after intraductal induction of *Ad-K8-Cre* adenovirus were prepared using the RNeasy kit (Qiagen, #73504) and subjected to microarray expression profiling at the Dana-Farber Cancer Institute Molecular Biology Core Facilities. The NuGEN Ovation V2 Amplification System was used to amplify and label samples. Biotinylated cDNAs were hybridized to MoGene-2_0-st Affymetrix Mouse Gene 2.0 ST Array [transcript (gene) version] chips. The raw CEL files were imported and processed with genepattern AffySTExpressionFileCreator tool (Broad Institute). Expression data was generated by AffySTExpressionFileCreator tool using the Robust Multi-array Average (RMA) algorithm as provided by the ’oligo’ package in Bioconductor. For mammary tumors, total RNAs were prepared from tumor aliquots previously snap-frozen and stored at -80°C by using the Zymo RNA extraction kit. RNA-seq for these tumor RNA samples were performed at the Dana-Farber Cancer Institute Molecular Biology Core Facilities by using an Illumina NovaSeq X Plus instrument. Visualization Pipeline for RNA-seq (Viper) analysis tool developed by Center for Functional Cancer Epigenetics (CFCE) at Dana-Farber Cancer Institute was used to generate standard outputs from RNA-seq data. Viper output data was normalized by using the DESeq2 module in GenePattern (https://www.genepattern.org/) for GSEA.

GSEA was conducted by using the standalone GSEA program (https://www.gsea-msigdb.org/gsea/index.jsp) and the Molecular Signatures Database (MSigDB) (https://www.gsea-msigdb.org/gsea/msigdb).

### Immunofluorescence

The dissected tumor tissues were fixed in 10% formalin and embedded in paraffin. Immunofluorescent labeling was carried out by following standard procedures, by incubating tumor tissue section with primary antibodies for ERα (MC-20) (Santa Cruz #SC542, 1:100), Phospho-STAT1 (Tyr701) (58D6) Rabbit Monoclonal Antibody (Cell Signaling Technology #9167T, 1:400), cytokeratin 8 (K8) (clone TROMA-1, Sigma-Aldrich # MABT329, 1:250), β-Catenin (BD Transduction Laboratories™ Purified Mouse Anti-β-Catenin, BD Biosciences #610154, 1:100), LEF1 (C12A5, Rabbit Monoclonal Antibody, Cell Signaling Technology #2230T, 1:400), and Ki-67 (D3B5, Rabbit Monoclonal Antibody, Cell Signaling Technology #9129T, 1:400), diluted using M.O.M.(TM) Basic Immunodetection Kit (Vector Laboratories # BMK-2202). The section was then washed with 1x PBS, and incubated with the secondary antibody [Goat anti-Rabbit IgG (H+L) Highly Cross-Adsorbed Secondary Antibody, Alexa Fluor 594 (Invitrogen A-11037), Goat anti-Rat IgG (H+L) Cross-Adsorbed Secondary Antibody, Alexa Fluo 647 (Invitrogen A-21247)] or [Goat anti-Mouse IgG (H+L) Cross-Adsorbed Secondary Antibody, Alexa Fluor™ 594 (Invitrogen A-11005), Goat anti-Rabbit IgG (H+L) Highly Cross-Adsorbed Secondary Antibody, Alexa Fluor™ Plus 647 (Invitrogen A32733TR)] for 30 min at room temperature.

## Statistics

All data are presented as mean ± SEM. Statistical comparisons between two groups were conducted with unpaired Student’s *t*-tests, whereas multi-group analyses were performed using two-way ANOVA followed by Tukey’s multiple comparison correction. Statistical significance was defined as \**p*≦0.05, \*\**p*≦0.01, and \*\*\**p*≦0.005.

## Data availability

Expression profiling datasets for premalignant MECs and tumors have been deposited to Gene Expression Omnibus (GEO) repository (https://www.ncbi.nlm.nih.gov/geo/) with the dataset identifiers GSE316272 and GSE316271, respectively.

## ACKNOWLEDGEMENTS

We thank Dr. Alan Cantor (Boston Children’s Hospital) for the *Runx1^L/L^* conditional knockout mouse line. This research was supported by NIH/NCI R21 grant (R21CA256468) and R01 grants (R01CA222560, R01CA295752) and by Brigham and Women’s Hospital Sundry Fund to ZL.

