## Supplementary Figures and Legends for "RUNX1-deficiency drives immune-active ER^+^ mammary tumorigenesis through activation of interferon signaling"

### Supplementary Figure Legends

**Supplementary Fig. 1:** Additional characterization of mutant MECs and mammary tumorigenesis in the indicated models. (a) Percentages of $YFP^+$ cells in mammary glands of indicated models at the last time point of chasing (i.e., 26 weeks or more). Statistics are based on comparison with percentages of $YFP^+$ cells from the induced *R26Y* control mice. ***$p \leqq 0.005$; ns: not significant. (b) Co-immunofluorescence for β-catenin and LEF1 in mammary tumors from the indicated models. Scale bars = 25μm.

**Supplementary Fig. 2:** FACS gating strategy for analyzing Sca-1 expression in either all luminal mammary epithelial cells (MECs) or YFP-marked luminal MECs.

**Supplementary Fig. 3:** Additional RNA-seq analysis of *RxPY*, *RbPY,* and *PRRY* mammary tumors. (a) Heatmaps showing high levels of expression of most genes in PRR-1 and/or PRb-6 tumors in mouse genomic regions spanning either *Yap1* or *Met* (highlighted). (b-c) GSEA data showing top enriched gene sets representing mouse mammary tumor (b) or human breast cancer (c) intrinsic subtypes extracted from Pfefferle et al (human intrinsic subtypes were based on the UNC308 data set and Combined855 data set)[1] in tumors with *Yap1* amplification (i.e., PRR-1 and PRb-6) in relation to other tumors. (d) Immunofluorescence for Ki67 in mammary tumors from the indicated models. Scale bar = 50μm.

**Supplementary Fig. 4:** FACS gating strategy for major immune cell populations in the tumor microenvironment. Representative plots for mammary tumors from each model are shown.

**Supplementary Fig. 5:** FACS gating strategy for major immune cell populations in the premalignant mammary gland (MG). Representative plots for mammary glands from each model are shown.

**Supplementary Fig. 6:** Additional analysis of immune-related gene and gene sets in RUNX1-deficient MECs. (a-d) GSEA data showing top enriched gene sets in the Hallmark collection from the MSigDB in *Runx1/Trp53*-null luminal MECs [compared to WT (a) or *Trp53*-null (b) or *Brca1/Trp53*-null (c) luminal MECs] or in MCF7 cells with *RUNX1* KD (d). Immune/IFN-related gene sets are highlighted. (e) UCSC genome browser view showing three RUNX1 binding peaks (arrows) in the *STAT1* gene (transcription direction from right to left); RUNX1 ChIP-seq data in MCF7 cells is based on GEO #GSE75070. (f) Negative correlation of levels of *RUNX1* and *STAT1* in human breast cancers, based on Breast Cancer Gene-Expression Miner v4.1 (bc-GenExMiner v4.1). (g-h) Kaplan-Meier curves showing the exact opposite correlations of *STAT1* (g) and *RUNX1* (h) levels with clinical outcomes, based on different subtypes of human breast cancers in K-M plotter (http://kmplot.com/analysis/).

**Supplementary Fig. 7:** Defining a *RUNX1*-low subset of $ER^+$ breast tumors from the TCGA cohort (cbioportal). (a) 91 and 74 cases from $ER^+$ breast tumor samples in the TCGA Pan-cancer cohort with the lowest and highest levels of *RUNX1* (see lower left plot) are assigned to the *RUNX1*-low and *RUNX1*-high groups, respectively. In this cohort, *RUNX1*-low tumors have slightly higher levels of *ESR1* expression (lower right plot). (b) Patient survival analysis showing *RUNX1*-low cases exhibited significantly worse overall survival. (c) *TP53* mutations are more significantly associated with *RUNX1*-low tumors. (d) GSEA data showing top enriched gene sets in the Hallmark collection from the MSigDB in *RUNX1*-low tumors (compared to *RUNX1*-high tumors); immune-related gene sets are highlighted. (e) GSEA plot showing a macrophage-related gene set in *RUNX1*-low tumors (compared to *RUNX1*-high tumors); gene set is derived from Nirmal et al [2]. (f) Significantly higher levels of *STAT1* expression in *RUNX1*-low tumors (compared to *RUNX1*-high tumors).

**a**

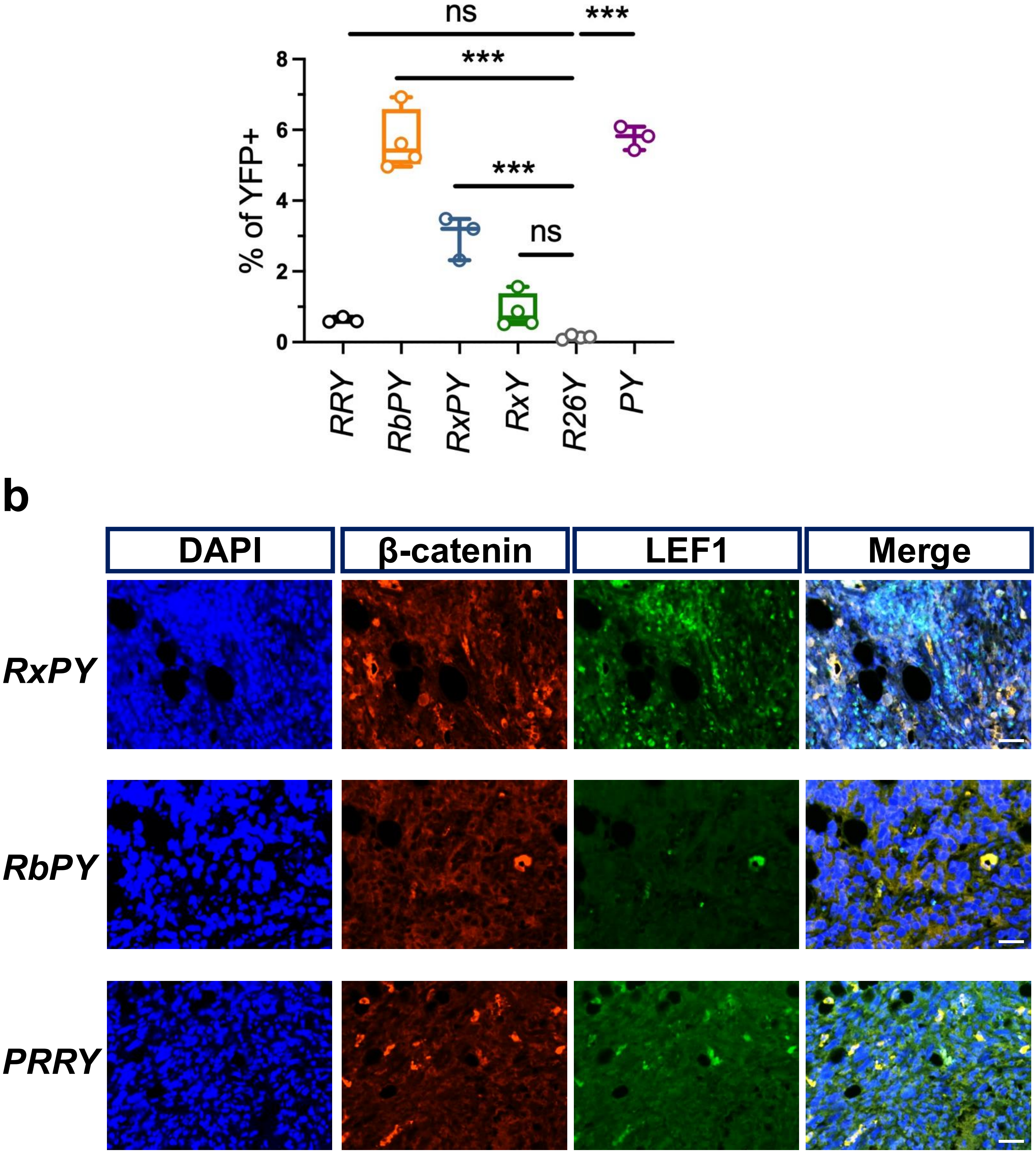
ns
***
***
***
ns
8
6
4
2
0
% of YFP+
RRY
RbPY
RxPY
RxY
R26Y
PY
b
DAPI
β-catenin
LEF1
Merge
RxPY
RbPY
PRRY

Supplementary Figure 2

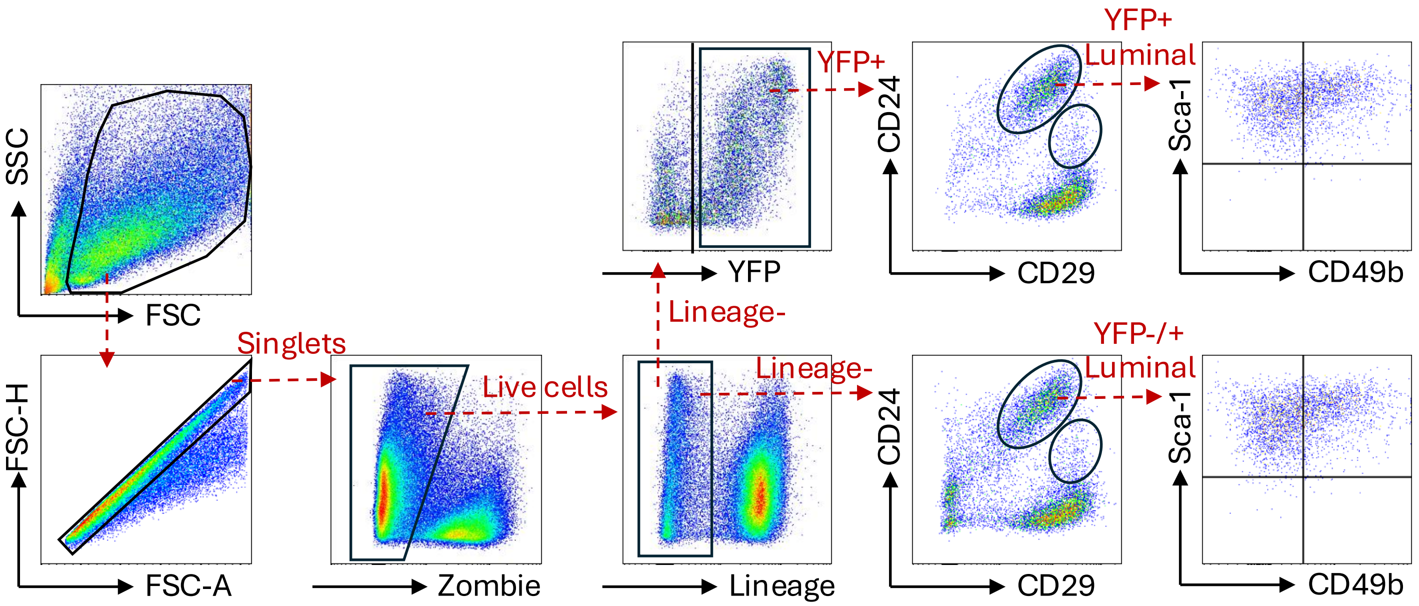

Supplementary Figure 3

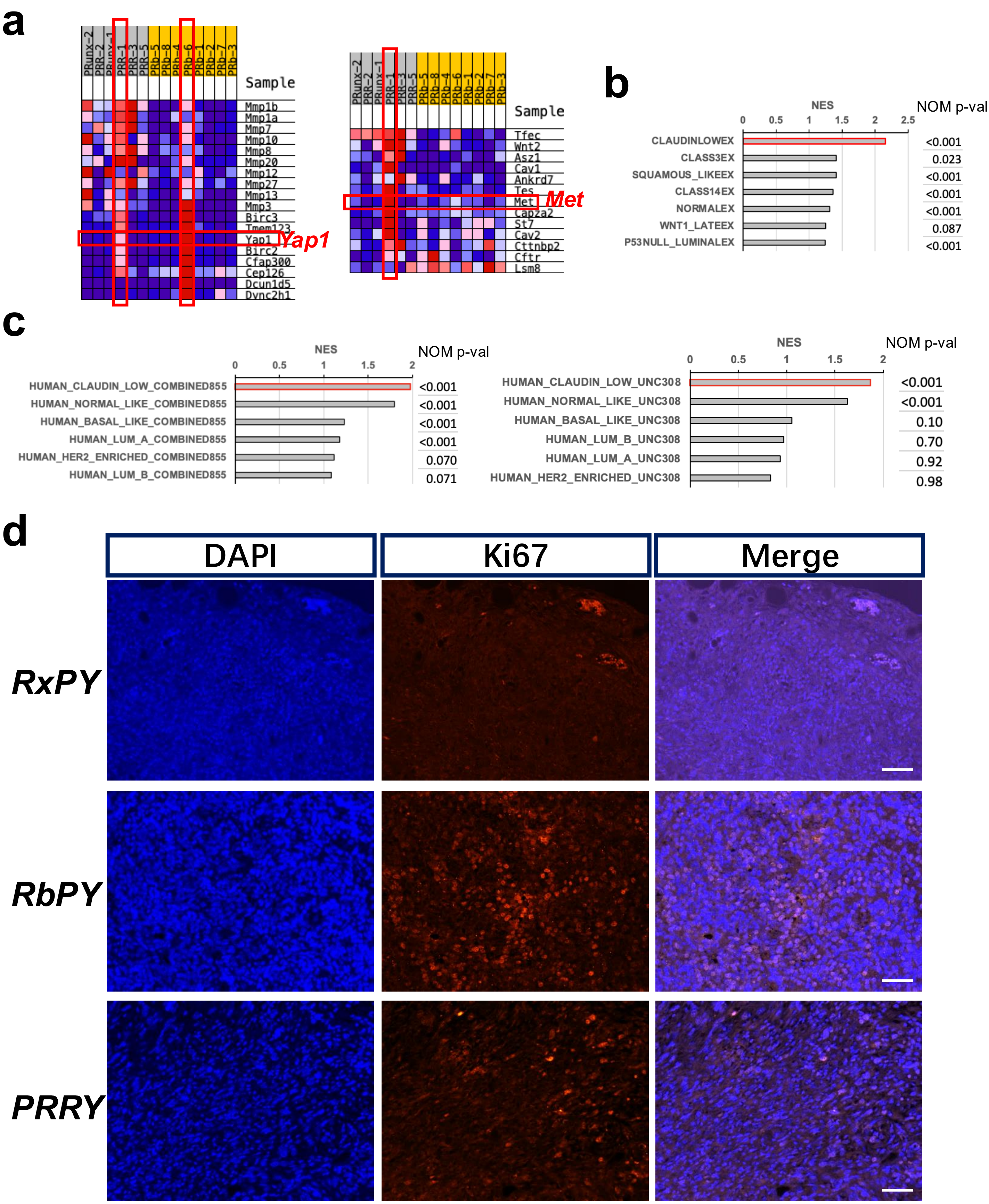

Supplementary Figure 4

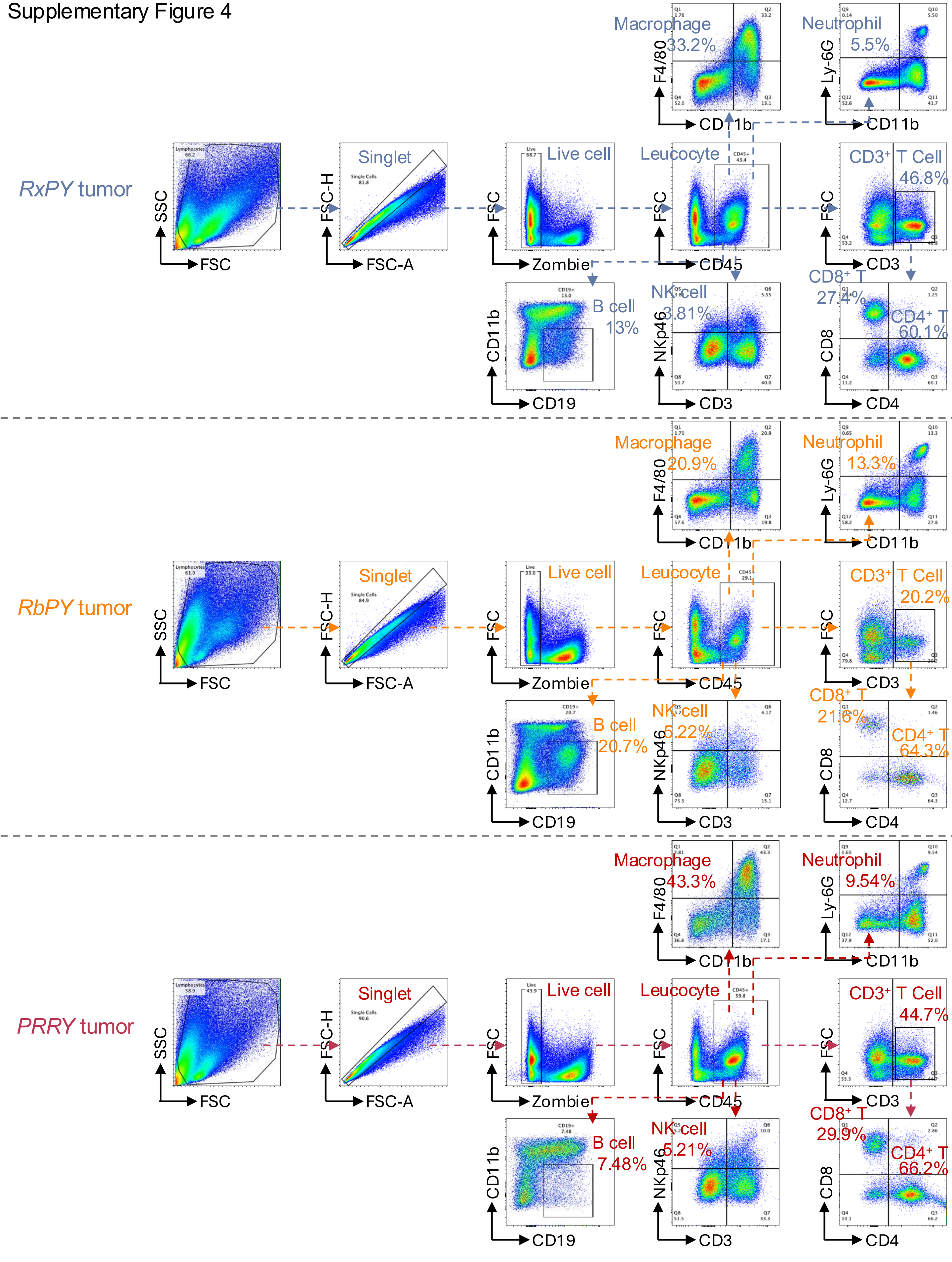

Supplementary Figure 5

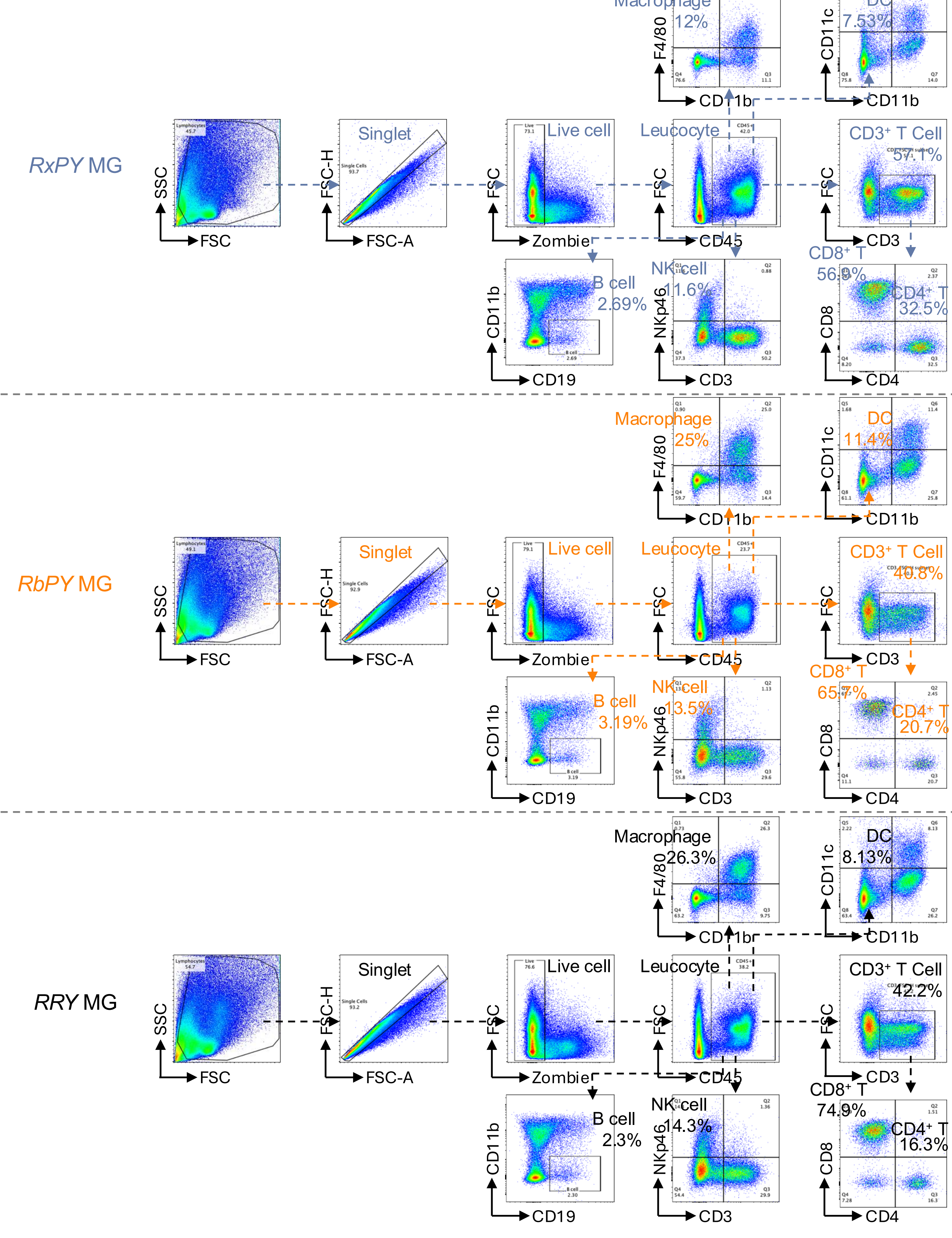

Supplementary Figure 6

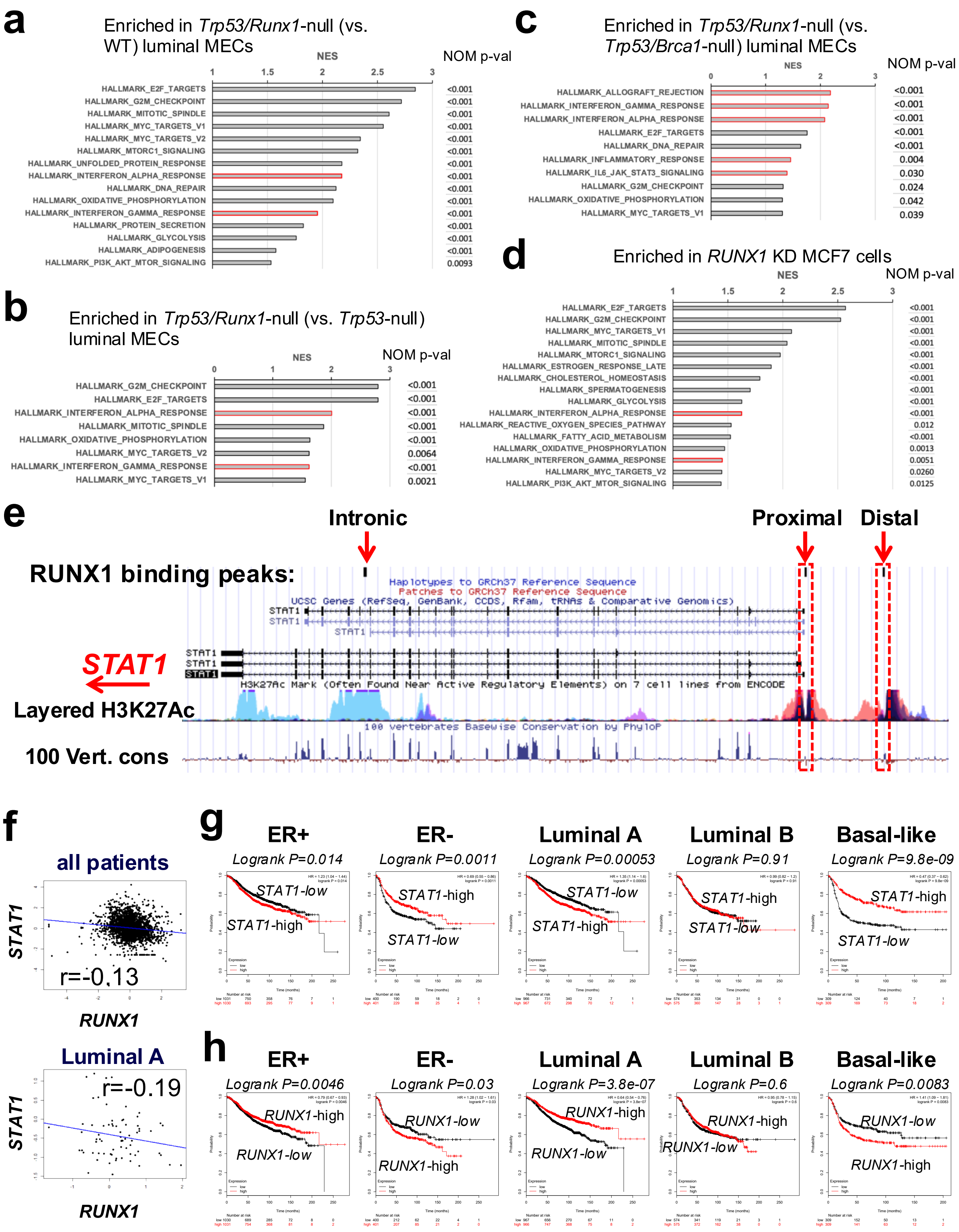

Supplementary Figure 7

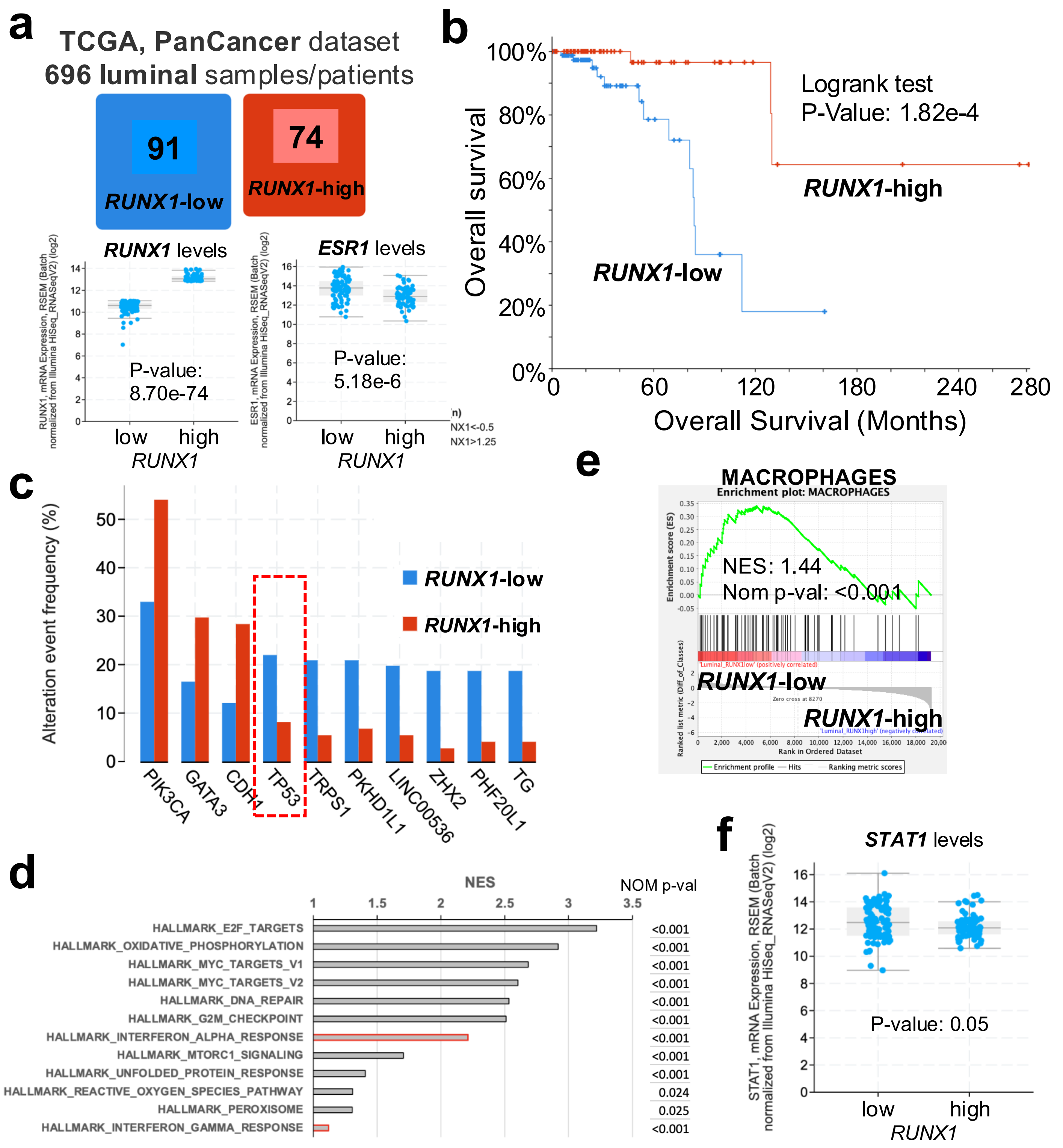
