## Supplementary Table 1 for "RUNX1-deficiency drives immune-active ER^+^ mammary tumorigenesis through activation of interferon signaling"

List of antibodies used in FACS analysis.

| REAGENT or RESOURCE | SOURCE | CLONE# | IDENTIFIER |
| --- | --- | --- | --- |
| Rat anti-mouse CD45 (BV605) | Biolegend | 3—F11 | Cat# 103155 |
| Rat anti-mouse TER-119 (BV605) | Biolegend | TER-119 | Cat# 116239 |
| Rat anti-mouse CD31 (BV605) | Biolegend | 390 | Cat# 102427 |
| Rat anti-mouse CD24 (PE) | Biolegend | M1/69 | Cat# 101808 |
| Hamster anti-mouse CD29 (APC) | Biolegend | HMβ1-1 | Cat# 102216 |
| Rat anti-mouse CD49b (PerCP/eF710) | eBioscience | DX5 | Cat# 46-5971-82 |
| Rat anti-mouse Sca-1 (PE/Cy7) | eBioscience | D7 | Cat# 25-5981-81 |
| Rat anti-mouse CD8a (BUV395) | BD | 53-6.7 | Cat# 565968 |
| Hamster anti-mouse CD3 (BUV737) | BD | 500A2 | Cat# 741716 |
| Rat anti-mouse CD4 (BV711) | Biolegend | RM4-5 | Cat# 100549 |
| Rat anti-mouse NKp46 (BV650) | Biolegend | 29A1.4 | Cat# 137635 |
| Rat anti-mouse CD45 (Alexa Fluor 700) | Biolegend | 30-F11 | Cat# 103128 |
| Rat anti-mouse F4/80 (BUV395) | BD | T45-2342 | Cat# 565614 |
| Rat anti-mouse CD19 (BUV737) | BD | 1D3 | Cat# 612782 |
| Rat anti-mouse CD11b (BV650) | Biolegend | M1/70 | Cat# 101239 |
| Hamster anti-mouse CD11c (BV510) | Biolegend | N418 | Cat# 117337 |
| Rat anti-mouse Ly-6G (BV421) | Biolegend | 1A8 | Cat# 127627 |
| Zombie NIR Fixable Viability Kit | Biolegend | N/A | Cat# 423106 |
| Zombie UV Fixable Viability Kit | Biolegend | N/A | Cat# 423108 |
